# Hybrid bioinks for embedded bioprinting of an artery model

**DOI:** 10.1101/2025.04.10.648100

**Authors:** Uxue Aizarna-Lopetegui, Ada Herrero-Ruiz, Monize Caiado-Decarli, Karla Chavarri, Lorenzo Moroni, Malou Henriksen-Lacey, Dorleta Jimenez de Aberasturi

## Abstract

The integration of biomaterials with living cells and stimuli-responsive materials can be employed to create bioinks capable of generating 3D *in vitro* models that better recapitulate native tissues. We introduce a multilayered artery model that combines hybrid multifunctional materials including a stimuli-responsive polymeric ink to mimic the *tunica adventitia* arterial wall, and an extracellular matrix (ECM)-based bioink for the *tunica media* artery layer. The stimuli-responsive hybrid layer integrates inorganic (plasmonic nanoparticles) and organic (polymers) components, providing structural support and introducing diverse functionalities to the system. The cell-laden bioink consists of human vascular smooth muscle cells (vSMC) within a hydrogel based on porcine artery-derived decellularized extracellular matrix (dECM) that fosters optimal cell growth and proliferation. An embedding bioprinting technique was employed for the fabrication of the multimaterial artery model consisting of concentric cylinders. The dimensions of the 3D model and the bioprinting parameters were fine-tuned to ensure effective crosslinking of the multiple concentric layers resulting in the creation of self-supporting constructs. We demonstrate the effectiveness of the hybrid bioink composition and bioprinting parameters in supporting cell viability and proliferation within the multilayered construct, expanding the possibilities of employing novel multi-component materials for the fabrication of 3D vasculature models resembling the structure of native blood vessels.

## 1. Introduction

There is a united interest in developing *in vitro* 3D models that can recapitulate the complex multi-tissue structure of human organs in healthy or pathological states. For example, for a correct fabrication of the blood vessel (whose composition and size varies enormously depending on factors such as subject age, health status, and location), the multi-tissue and multilayered structure of vasculature must be considered. In general, human arteries consist of three main tissue layers: the outer *tunica adventitia*, the middle *tunica media*, and the inner *tunica intima*, the latter surrounding the vessel lumen and lying in direct contact with the bloodstream.^[1,2]^ Each tissue layer has a particular cellular and non-cellular composition based on function and geometry, with a clear distinction made between large vessels (> 6 mm in diameter) or small vessels (< 6 mm in diameter).^[3]^ A variety of techniques including melt electrowriting and electrospinning,^[4–6]^ decellularization of xenogeneic vessels,^[7]^ tubular casting,^[8]^ and ice templating^[9]^ approaches have been described for the fabrication of blood vessel-like structures. In these models, diverse natural physiological and pathological attributes, including size, geometry, and physico-mechanical properties, have been replicated. However, reconstructing with anatomical accuracy the intricate hierarchical structure of the blood vessel wall, i.e. ensuring uniform cell distribution, proper cell morphology, and appropriate tissue cohesion to achieve functional tissue, remains a significant challenge.^[10]^ To overcome these limitations, bioprinting has emerged as a powerful tool for developing structurally complex and multilayered constructs for tissue modeling.^[11–13]^ Extrusion-based bioprinting, the most widely used method for the layer-by-layer creation of self-supporting blood vessel models, is however still unable to replicate the intricate layered structure of native vessels. This limitation arises from the difficulty in constructing self-standing structures with multiple concentric layers using low-viscosity gels that adequately support cell growth and spreading.

While extrusion-based 3D printing has been used to create single-layered vascular analogs with embedded smooth muscle cells and seeded endothelial cells to model thromboinflammatory responses,^[14]^ as also previously reported by some of the authors for the bioprinting of horizontally layered simplified artery models, ^[15]^ these capabilities have not yet been extended to multilayered bioprinting of tubular structures. To successfully incorporate the relevant cell types into tall and layered cylindrical geometries, new bioprinting approaches are being designed and developed.^[16–18]^ For example, to achieve multi-layered vessels, various groups have explored the use of coaxial-extrusion bioprinting where a concentrically assembled nozzle is employed to form perfusable vessels.^[19]^ Considering that the formation of cylindrical structures via extrusion bioprinting involves the deposition of material in a vertical geometry, as the height of the structure increases, so does its center of mass, consequently increasing the risk of structural failure or collapse. Achieving self-supporting multilayered structures with adequate structural integrity requires concessions in the use of high-viscosity inks, potentially compromising cell viability in the process.^[20,21]^

Nevertheless, low-viscosity inks can be bioprinted by the use of sacrificial support baths, which enable the creation of vascularized and more complex organs, including full-size models based on low-viscosity cell-friendly bioinks.^[22–24]^ These approaches not only overcome the deformation and collapse experienced by low-viscosity bioinks upon extrusion, but also provide the necessary physical support for crosslinking to take place, while allowing the construction of stable and freeform structures, including blood vessel bifurcation. Since its introduction by the use of Pluronic F-127, various materials including natural polymers such as gelatin,^[25]^ agarose,^[26]^ xanthan gum,^[27]^ or hyaluronic acid,^[28]^ as well as cell-laden extracellular matrixes (ECMs)^[29]^ and functional cell containing tissues,^[30]^ have been described as suspension baths for embedding bioprinting. These support materials should exhibit gel-like properties in the absence of stress, but transition to a fluid state when stress surpasses the material’s yield stress, enabling the nozzle or needle to move freely while extruding inks. In addition, the support material should be self-healing, regaining its microstructure and maintaining the printing fidelity of the deposited ink, after the nozzle or needle has passed.^[31,32]^ Finally, in the case of cellularized structures, the support material itself and components or techniques used in its removal or degradation must be biocompatible. To the best of our knowledge, there are no reports on the use of multi-layered bioinks for the development of multilayered *in vitro* blood vessels by bioprinting to study vascular physiological and pathological processes.

Aside from the engineering of support materials, remarkable progress is being made in the design and development of bioinks for the fabrication of biological constructs via bioprinting.^[33–36]^ Traditionally, single-component natural and synthetic hydrogels have been used to mimic tissue ECMs, but they often yield mixed results due to their suboptimal mechanical and biochemical properties.^[37]^ In contrast, blended compositions or naturally heterogeneous (i.e decellularized ECMs) biomaterials offer more positive outcomes by overcoming these limitations and providing a more complete representation and functionality of the ECM.^[38]^

One novel method to gain mechanical and biochemical versatility involves the fabrication of hybrid materials with multifunctionality. The incorporation of inorganic nanoparticles (NPs), which act as stimuli generators^[39]^, into organic polymeric matrices can be exploited for the generation of stimuli-responsive hybrid materials that can be precisely tuned to mimic diverse processes of biological significance.^[40]^ Harnessing the potential to manipulate and regulate tissue and organ behavior is driving the trend towards 4D printing, holding promise for creating mature, functional tissue constructs which better mimic the biological and anatomic attributes of human organs.^[41]^ The creation of bioinks based on advanced stimuli–responsive materials, engineered to dynamically respond to external chemical (e.g. humidity, pH) and physical (e.g. temperature, electric, magnetic, or acoustic fields) stimuli, holds considerable promise in generating multifunctional blood vessel models.^[42]^ Such “smart properties” are generally achieved via the addition of gold,^[43,44]^ silver,^[45]^ iron oxide,^[46]^ and piezoelectric,^[47]^ NPs, and carbon nanotubes, into polymeric matrices.^[48]^ For example, the combination of thermoresponsive polymers with gold NPs that can generate heat upon laser irradiation,^[49,50]^ iron oxide NPs that present magnetic properties and can induce polymer swelling,^[51]^ or piezoelectric NPs that can generate electric potentials.^[52]^ The combination of stimuli-responsive materials with cell-laden bioinks containing essential ECM components, such as those facilitating high biomimicry and cell compatibility, holds promise to recreate multi-tissue organ models. To improve the biological fingerprint of cell-laden bioinks and better mimic the native tissue environment, the focus has turned to the use of dECM.^[53]^ Obtained by removing all cellular components from the tissue source, dECMs offer an excellent matrix for cellular functions including structural support, cell adhesion, differentiation, signaling, and survival, as well as regulation of protein complexes and growth factors.^[54]^

Herein we present the development of a self-standing model comprised of concentric cylinders of stimuli-responsive and dECM-based layers, mimicking the structure of blood vessels. We demonstrate the feasibility of using embedding bioprinting to create a concentric multimaterial artery model, developing two innovative inks for the *tunica adventitia* and *tunica media* layers. Whilst the elastic and thermoresponsive outer layer provides structural support, the inner dECM-based layer embedded with smooth muscle cells creates a biocompatible environment for cell growth and spreading. These findings describe the potential of multimaterial hybrid bioinks for bioprinting applications, specifically for the development of *in vitro* models suitable to study blood vessel mechanobiology.

## 2. Results and Discussion

### 2.1. Development and characterization of inks for a 3D bioprinted multilayered vasculature model

For the fabrication of the multilayered cylindrical artery model, we employed an elastic and stimuli-responsive polymeric ink for the *tunica adventitia*, and a dECM-based bioink with embedded smooth muscle cells (vSMCs) as the *tunica media*. To enhance structural integrity and crosslinking between the *tunica adventitia* and *tunica media*, we incorporated gelatin methacryloyl (GelMA) into both inks, thus serving as an interfacial material to facilitate the attachment between the two layers. GelMA offers various advantages such as allowing the simultaneous photo-crosslinking of both layers upon bioprinting, offering customizable mechanical properties, and improving biocompatibility. Considering that collagen is the principal component of the blood vessel wall, and that GelMA is a chemical derivative of collagen, inclusion of GelMA was biologically justifiable.

#### 2.1.1. Stimuli-responsive ink

The *tunica adventitia*, composed of connective tissue, plays a critical role in providing structural support. To accurately replicate this layer, we developed an elastic and thermoresponsive ink, referred to as “hybrid ink” or “hybrid gel” depending on its unphotopolymerized or photocrosslinked state, respectively, exhibiting tunable mechanical properties in response to temperature changes. We based this hydrogel formulation on a previously described thermoresponsive ink, which showed light-sensitive stimuli responsiveness thanks to the incorporation of AuNPs embedded in thermoresponsive polymeric networks.^[15]^ We showed that NIR-induced pulsed irradiation led to thermoresponsive gel contraction, which induced the expression of mesenchymal activation gene signatures associated with the YAP/TAZ pathway, central to the mechanotransduction network. To make self-standing multilayered concentric cylinder constructs, the stimuli-responsive ink was modified and optimized (for full details, see Experimental Section). The final composition of the new stimuli-responsive polymeric ink consisted of a polymeric blend comprising 16 % (w/v) N-isopropylacrylamide (NIPAm), 6 % (w/v) poly(ethylene glycol) diacrylate (PEGDA), 2 % (v/v) poly(ethylene glycol)dimethacrylate (PEGDMA), 5 % (w/v) gelatin methacryloyl (GelMA), 0.5 mM AuNRs, and 0.1 % (w/v) lithium phenyl-2,4,6-trimethylbenzoylphosphinate (LAP) (**Figure 1A**). The inclusion of PEGDA (700 Da) allowed us to adjust the lower critical solution temperature (LCST) of the thermoresponsive polymer NIPAm, bringing it closer to physiological body temperature. PEGDMA (Mn = 550) and GelMA, in contrast, were included to improve elasticity and structural crosslinking, respectively, using LAP as the photo-polymerization photoinitiator. As expected, the inclusion of PEGDMA resulted in a decrease in the Younǵs Modulus (a measure of material tensile strength), in agreement with values for native arteries reported in literature (**Figure 1B**).^[55]^ Compression and tensile stress-strain tests revealed that the elongation of the material at break was higher in the PEGDMA-containing inks (25.8 % ± 4.5) compared to those without (11.0 % ± 3.1), indicating an increase in the elasticity.

**Figure 1.**
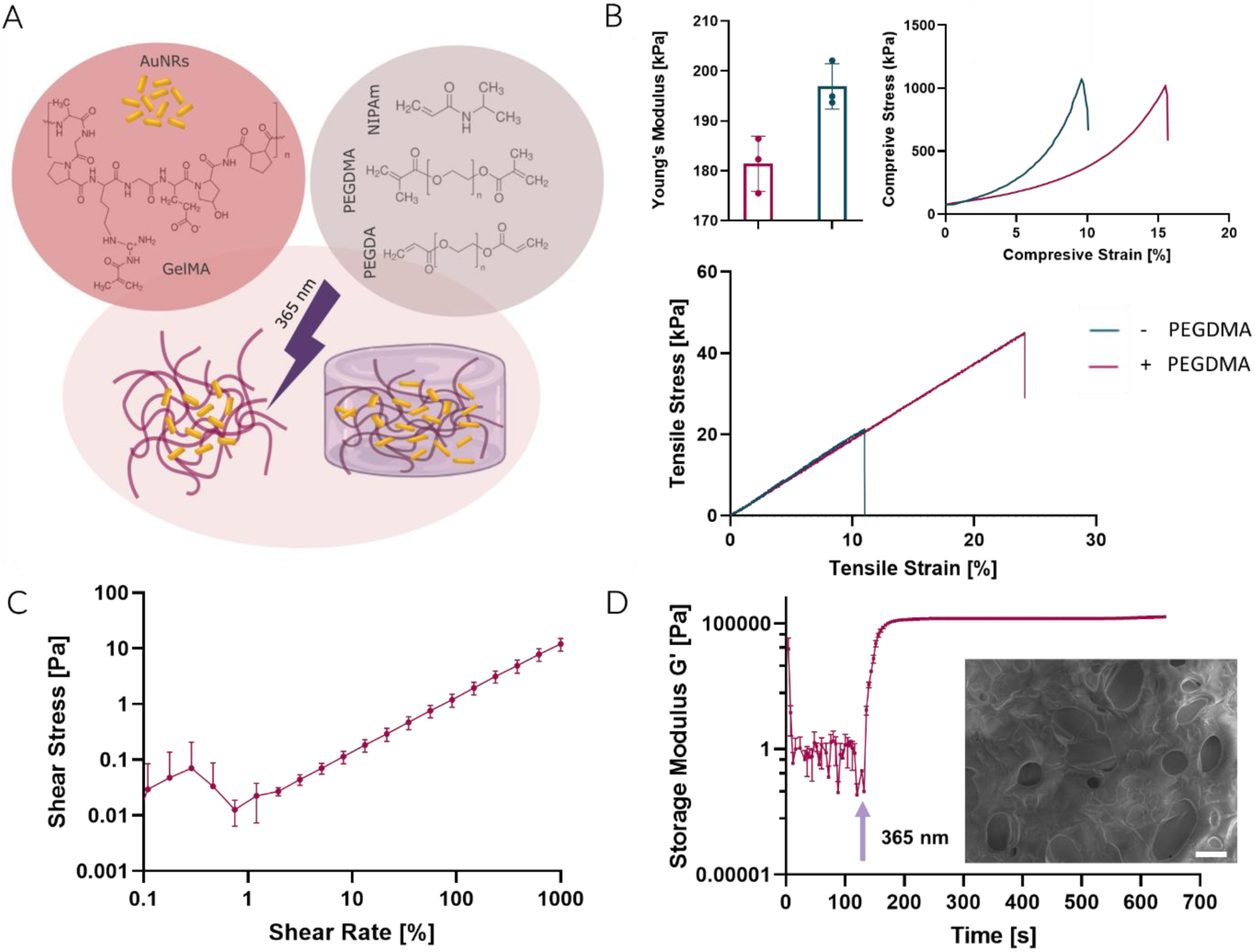
Hybrid ink development and characterization. A) Schematic representation of the ink preparation and its photo-crosslinking upon UV irradiation for the formation of stable covalently crosslinked polymeric networks. B) Mechanical characterization showing the differences in Young’s modulus, compression, and tensile strain tests of the hybrid gel, with and without PEGDMA. C) Rheological characterization of the final hybrid ink composition including the flow curve at varying shear rates, and (D) thixotropy test combined with *in situ* photo-crosslinking (mean ± SD, n=3) and a SEM micrograph image of the hybrid gel (SEM insert: scale bar 20 µm).

To achieve maximum AuNR heating upon light irradiation, AuNRs were synthesized with an LSPR in the NIR, in resonance with the employed 808 nm laser. In agreement with previous findings,^[15]^ we observed a slight red-shift of the LSPR of gel-incorporated AuNRs due to the higher refractive index of the hydrogel (**Figure S1**). The thermoresponsive nature of the hydrogel was characterized by differential scanning calorimetry (DSC), where the LCST was observed at 36 °C (**Figure S2**), and further confirmed by the increase in storage modulus observed around the hydrogel transition temperature (**Figure S3**), in agreement with previously reported findings.^[56]^ We employed a variable bandwidth laser to measure the *in situ* heating capacity of the AuNRs embedded in the hybrid gel. In preliminary experiments we observed maximum heating using a bandpass filter with a bandwidth of 730 – 850 nm, encompassing the maximum absorbance (LCST) of the AuNR (**Figure S4A**). By employing a pulse generator, we successfully controlled hydrogel heating around its LCST in a rapid and cyclic manner (**Figure S4B**). These results confirm the photothermal conversion ability of AuNRs, even when immobilized in the hybrid gel. Importantly, irradiation had no irreversible effect on the polymer morphology and stability, confirming the non-invasive character of this technique to achieve hydrogel expansion and contraction, highlighting its potential for pulsed heating applications. We applied rheological measurements to determine the shear thinning behavior, observing an increase in shear stress with increasing shear rate, which supported the use of the hybrid gel for extrusion-based 3D printing (**Figure 1C**). To simulate polymerization post-printing, we exposed the material to UV irradiation for 30 seconds, observing the *in situ* changes in storage modulus over time. The ink maintained a liquid state, where the loss modulus (G’’) remained higher than the storage modulus (G’) until it underwent near-instantaneous crosslinking upon exposure to UV light. At this stage, G’ surpassed G’’, confirming the desired photo-rheological properties of the formulation (**Figure 1D, Figure S5A**). At this point it is important to note that before being crosslinked upon UV exposure, the material presents low viscosity for conventional extrusion-based bioprinting approaches. Notably, the crosslinked material exhibited a typical gel behavior at 37 °C, a temperature at which the 3D models were kept favoring cell culture conditions, evidenced by the frequency and amplitude sweeps performed at the target temperature (**Figure S5B-C**). The modulus of the material remained constant in time at a value around 100 kPa. In particular, the material showed a typical yield stress-like response at 37 °C and low applied strains, with a dominant (elastic) solid-like response characteristic of hydrogel-based materials. The elastic component (G’) consistently dominated over the viscous part (G’’), with both moduli remaining independent of the applied frequency at low deformations. The material presented a linear viscoelastic (LVE) response until starting to yield at high shear stresses, where a deformation-driven solid-to-fluid transformation was observed. SEM imaging revealed a polymeric network with pores of < 50 µm in diameter, ideal for the facile diffusion of oxygen and nutrients to the cell-containing *tunica media* (**Figure 1D**).

#### 2.1.2. Cell-laden bioink

The *tunica media*, primarily composed of human vascular smooth muscle cells (vSMCs), elastin, and collagen fibers, confers mechanical strength and enables the dilation and contraction of arteries. To establish a biologically relevant ECM microenvironment, we developed a bioink composed of decellularized porcine pulmonary artery tissue. This choice was based on previous reports on its similarity in ECM composition to human blood vessels,^[2]^ biological compatibility with human vSMCs,^[57,58]^ and its relative abundance. To remove the cellular component of the extracted tissue, while maintaining specific matrix components, scaffolding elements, and cell binding motifs for cell growth and proliferation, a protocol for thick-walled tissue decellularization based on the chemical and enzymatic removal of cells was adapted from the work of Koenig et al., as detailed in **Table S1** and **Figure S6**.^[59]^ Histological analysis were conducted to assess the effectiveness of the decellularization process, verifying the retention of the most relevant structural components of native arteries, and the absence of cells, as measured by hematoxylin and eosin, DAPI staining and DNA measurements (**Figure 2A, Figure S7A**).^[60]^ The dECM retained structural proteins such as collagenous fibers, confirmed using polarized light and Picrosirius Red staining (**Figure 2B, Figure S7B**).^[61]^ Image analysis of the histological sections of native and decellularized ECM showed a loss of 15 % of collagen (all types) during the decellularization process (for full details, see Experimental Section). Additionally, elastic fibers were visualized through Verhoeff Van Giesson staining of the histological sections (**Figure 2C, Figure S7C**). The presence of a fibrous surface, in both native and decellularized ECM, could also be observed via SEM (**Figure S7D**).

**Figure 2.**
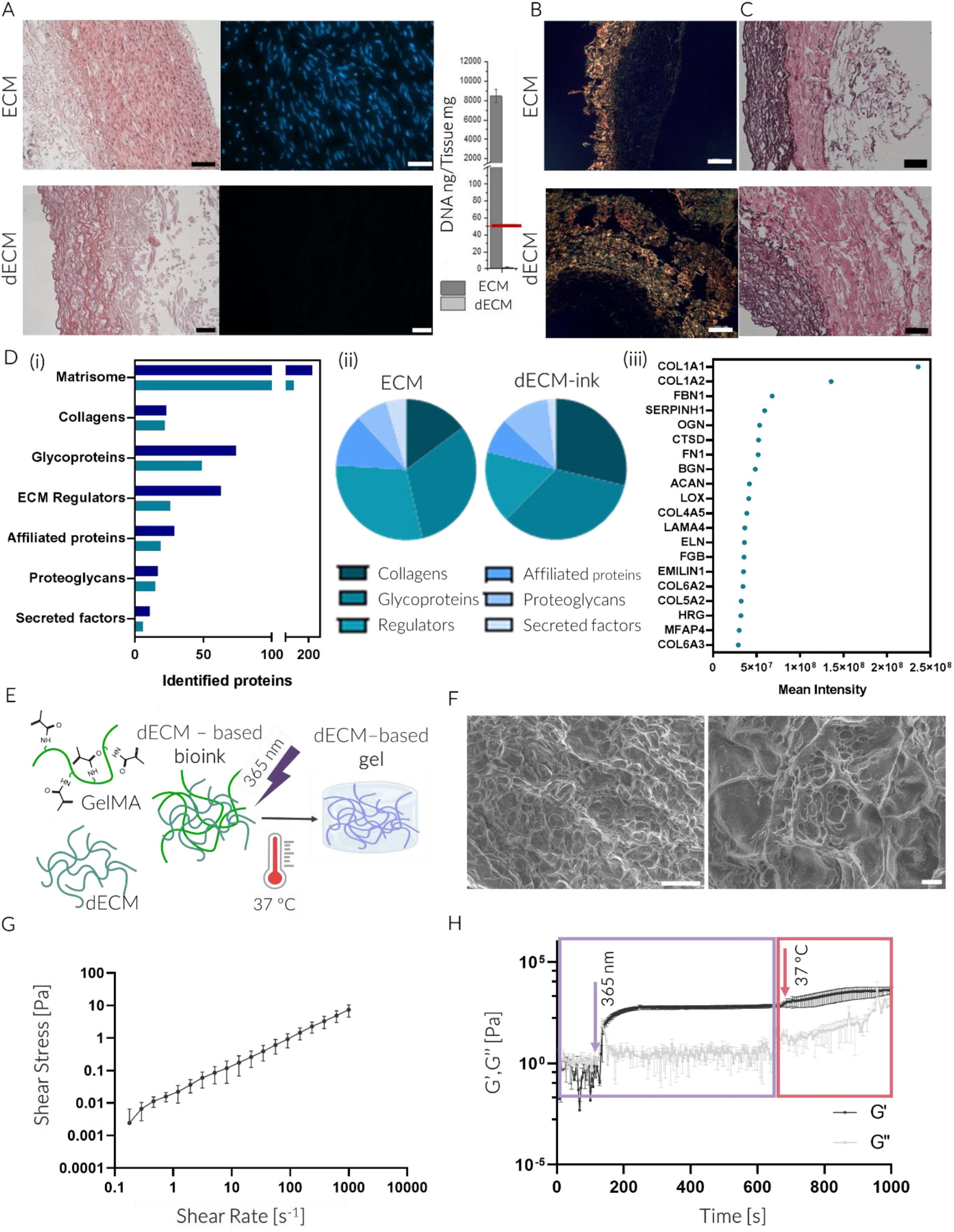
Development and characterization of the dECM-based bioink. A) Characterization of the decellularization process of porcine pulmonary artery including representative images of Hematoxylin and Eosin (H&E) and DAPI fluorescence imaging of native (ECM) and decellularized (dECM), and DNA quantification of pulmonary artery tissue (scale bars: H&E, 100 µm; DAPI, 50 µm). B-C) Evaluation of the structure and composition of the ECM and dECM including the study of the presence of collagenous fibers by polarized light of Picrosirius Red staining (scale bars: 200 µm) (B) and Verhoeff Van Giesson stain showing elastic fibers of the histological sections (scale bars: 100 µm) (C). D) Proteomic analysis of ECM and dECM-ink including (i) the quantification of identified matrisome related proteins, (ii) pie charts of the relative intensity of the matrisome categories for the ECM and dECM-ink, and (iii) the 20 most abundant genes associated with the dECM-ink, based on the mean intensity of the proteomic analysis. E) Schematic representation of the generation of the dECM-based gel through thermal and light-based crosslinking. F) SEM images of the crosslinked hydrogels containing 3 w/v % GelMA and 0.1 w/v % dECM (scale bars: left, 100 µm; right, 20 µm). G-H) Rheological properties of the dECM-based bioink through flow curve (G) and viscoelastic properties of the material upon crosslinking (H) including photo-crosslinking (purple) and thermal (red) crosslinking mechanisms (mean ± SD, n=3).

The porcine pulmonary artery dECM was subsequently pulverized to a powder, digested under acidic conditions, and the pH neutralized. We conducted a detailed proteomic analysis of this dECM bioink and its native ECM, analyzing their composition on various levels (see SI for full details) ^[62]^ Firstly, we deduced the proportion of proteins homologous to the human species (97 %), and focusing on these proteins of translation interest, we thus subjected our analysis specifically to those of the matrisome (**Figure 2Di-ii, Figure S8**). The matrisome, considered to include all ECM proteins and associated molecules that form a structural and functional network surrounding cells, is of primary importance in understanding the biological composition of the ECM and dECM bioink. Our findings, whilst extensive, highlighted: a) the high reproducibility of the decellularization process, as shown by principal component analysis (PCA) clustering (**Figure S8D**), in conserving matrisome-associated proteins; b) a loss of approximately 37 % of matrisome proteins upon decellularization; and c) retention of collagen gene and protein expression during the decellularization process (23 vs 22 collagen proteins in ECM and dECM, respectively, corresponding to 10 and 16 % of the total matrisome) (**Figure S9**). This latter finding is consistent with the collagen quantification analysis of histological sections described above (**Figure 2B, Figure S7B**). The increased resistance of collagens to decellularization and digestion procedures can be attributed to their primary role in providing structural support, which naturally contributes to their greater durability. Interestingly, when evaluating the genes associated to the proteins that presented the 20 highest mean intensity counts in the ECM and dECM-ink (**Figure 2Diii** and **S10**), in both cases the genes with the highest intensities were collagens: COL6A3 for the ECM, and COL1A1 for the dECM-ink. Among the proteins identified in the dECM-ink, COL1A1 and COL1A2 were the most abundant, followed by Fibrillin-1 (FBN1) and Heat Shock Protein 47 (SERPINH1). To assess how the ECM might affect cellular functions, we conducted enrichment analyses, a statistical approach used to determine if specific gene sets, proteins, or other biological entities are significantly overrepresented or underrepresented in a dataset compared to what would be expected by chance. This method uncovers the biological processes, functions, or pathways associated with the data under study (full details and results are shown in the SI). As can be observed in **Figure S11 and Figure S12**, whilst genes encoding proteins related to coagulation, hemostasis, and fibrinolysis were observed with highest presence, in the case of the dECM ink, genes related to the organization and maintenance of the ECM, including supramolecular fibre and collagen fibril organization, followed by elastic fibre assembly were most predominantly found.

We studied the dECM thermal gelling ability between 30 – 50 °C, observing an increase in the gel modulus due to self-assembly of collagen fibers (**Figure S13A**).^[63,64]^ In proof-of-concept experiments, we explored the possibility of chemically gelling the dECM using ruthenium and sodium persulfate combined with visible light photo-crosslinking agents. This method triggers the conversion of tyrosine groups, commonly present in collagen, into di-tyrosine leading to the formation of covalent crosslinks within the matrix, as previously reported.^[65]^ We observed successful gel formation upon irradiation with 405 nm light (**Figure S13B**), supporting the potential of using porcine artery dECM as a stand-alone gel with biocompatible crosslinking mechanisms (heat and light). However, considering our *tunica adventitia* gel lacks dECM, and previous findings had discussed the importance of achieving cross-linking between adjacent materials,^[56]^ we chose to adapt the *tunica media* bioink by including GelMA, thus achieving thermal and chemical crosslinking due to the collagen component and LAP, respectively (**Figure 2E**). An in-depth study on the biocompatibility of the *tunica media* dECM-based bioinks with human vSMCs, specifically pulmonary artery smooth muscle cells (hPASMCs), was conducted. Firstly, we assessed potential cytotoxicity due to soluble components leaching from gelled dECM and GelMA bioinks. Excellent vSMCs biocompatibility was observed after exposure to gelled bioinks (**Figure S14A**). We thus embedded vSMCs at a final concentration of 1.5·10^6^ cell/mL, in dECM and GelMA bioinks of various polymeric concentrations to further analyze cell viability and morphology over time. We observed that at dECM concentrations above 0.1 % (w/v), the cell viability decreased significantly, which may be related to the increase in substrate stiffness (**Figure S14B**). In the case of GelMA hydrogels, ≤ 1 % (w/v) resulted in inadequate gelling and the 3D nature of the gel was lost allowing vSMCs to adhere to the underlying substrate (**Figure S14C**). At higher GelMA concentrations, gelation was successful but the balance between GelMA concentration and expected cell morphology (i.e. cell spreading) was affected, with concentrations ≥ 5 % (w/v) affecting cell viability. In mixed dECM/GelMA formulations, we fixed the dECM concentration to 0.1 % (w/v) and studied GelMA concentrations of 2.5, 3, and 5 % (w/v) (**Figure S14D**). We observed little differences in the cell viability and/or morphology, and thus chose an intermediate value of 3 % GelMA, 0.1 % dECM with 0.1 % LAP (all w/v). SEM imaging revealed that this dECM-based bioink presented a fibrous surface with void areas of an average size of <10 µm, compatible with cellularization (**Figure 2F**).

The rheological properties of the selected dECM-based bioink and of the final crosslinked gel were evaluated to determine its suitability for bioprinting. The material exhibited shear-thinning behavior, essential for bioprinting (**Figure 2G**). The dual crosslinking of the material was studied by the *in situ* photo-crosslinking of the material and followed by an isothermal process at 37 °C (**Figure 2H**). The bioink remained in liquid state (G’’>G’) until crosslinked by UV irradiation. The modulus of the gel further increased upon raising the temperature to 37 °C, in agreement with the thermal gelation of the dECM. The final formulation in a gel form, as opposed to the liquid ink, showed a solid viscoelastic behavior at 37 °C, where the storage (G’) was higher than the loss (G’’) moduli and both were parallel and independent of the applied amplitude or frequency (**Figure S15**).

### 2.2. 3D bioprinting of multilayered vasculature model

Having shown that the chosen material for the *tunica adventitia* layer could recapitulate the mechanical properties of arterial wall ECM and that the cell-laden bioink provided a suitable environment for vSMCs to expand and proliferate, the materials were 3D bioprinted. Initial trials involved extruding the materials individually or printing them simultaneously to generate concentric cylinders. Regardless of the high post-printing cell viability, the dECM-based composition exhibited no structural fidelity upon bioprinting at room temperature (RT) (**Figure S16A**). Similarly, the hybrid ink collapsed when extruded at RT; however, by decreasing the printing temperature to 5 °C, 1 cm tall self-standing cylinders were obtained (**Figure S16B**). Despite aiming to enhance viscosity by printing the materials at 5 °C, limitations in the bioprinting setup allowed the cooling unit to be applied to only one material at a time. Thus, when moving to the simultaneous bioprinting of the materials for the generation of concentric cylinders, bioprinting fidelity was only obtained if the polymeric concentration was increased (**Figure S17**). As This environment does not greatly support cell growth and proliferation, we focused on identifying a bioprinting technique supportive of maintaining structural fidelity and cell viability. In this regard, the use of a support baths, e.g. gelatin or agarose-based baths, and the recently published *CLADDING* bioprinting^[32]^ method, were investigated. A coacervation process and a variety of washing approaches were employed to optimize the preparation of supporting baths (for full details, see Experimental Section). We investigated material transparency at 365 nm (the wavelength employed for the crosslinking of the constructs), and material stability at 37 °C, to determine the ideal support bath material to bioprint bi-layered cylinders of 1 cm in height. Whilst the gelatin-gum arabic-based bath had optimum transparency, it proved incompatible with the dECM crosslinking, which occurs at 37 °C (**Figure S18, S19A**). Similarly, whilst the agarose bath was thermally stable at 37 °C, its poor transparency required excessive crosslinking times, which were ultimately inefficient and resulted in high cell death (**Figure S19B**). *CLADDING* bioprinting provided a bath that offered improved transparency at 365 nm and sufficient stability at 37 °C to allow the thermal crosslinking of the dECM component before the slurry melted, after washing with PBS to effectively compact the slurry gel microparticles (**Figure S18)**. The optimal duration of printing and curing within the support bath is crucial for preserving cell viability in the constructs. According to Bessler et al., viability decreased with longer incubation times: 80% at 60 minutes, 70% at 90 minutes, and 50% at 120 minutes.^[66]^ We observed that starting the washing of the bath and supplying the tissue analogs with media after 45 minutes of incubation at 37 °C, sufficient to ensure the thermal crosslinking of the dECM component and melting of the support, was optimal in terms of maintaining high levels of cell viability (**Figure S20**).

#### 2.2.1. Embedded 3D bioprinting of single-component cylinders

In order to optimize the bioprinting and crosslinking parameters for each ink, we first explored the effect of bioprinting pressure, feed rate, layer thickness, angle between the consecutive layers, and nozzle diameter, obtaining self-standing cylinders, printed within the support bath, of separate bioinks (**Table S2**). To achieve superior shape fidelity and support thin structures avoiding collapse, 25G metallic needles measuring 2.54 cm in length were selected for the embedded bioprinting of both inks. Once deposited, the structure was crosslinked for 30 s using a 365 nm LED source. With regards to the hybrid ink, 1 cm tall self-standing cylinders with high printing fidelity were achieved (**Figure 3A**). We employed multiphoton confocal microscopy to characterize the cylinder cross-section, taking advantage of the embedded AuNRs which offer strong two-photon excited photoluminescence (2PEL) signals upon irradiation (**Figure 3B**).^[67]^ This technique allows us to achieve exceptionally high levels of resolution in the XYZ plane without the use of additional fluorescent markers to label the material. With regards to the dECM-based bioink cylinder, we successfully bioprinted cell-laden cylinders, but due to the contractive nature of hPASMCs^[68,69]^ when interacting with the biocompatible matrix which was supportive to cell growth, we observed contraction or collapse of the cylinder 24 hours post removal of the bath (**Figure 3C, Figure S21A).** Whilst the integration of support baths has significantly broadened the capabilities of extrusion hydrogel bioprinting, it has simultaneously introduced complexities and interactions that remain incompletely understood. Of particular importance is the interaction between the bioink and support bath materials themselves, pre- and post-crosslinking, both of which are hydrogel networks in a state of prolonged contact.^[70]^ Several studies describe that osmotic pressure gradients facilitate solvent exchange between the bath and ink (pre-/post-crosslinking), potentially altering concentration-dependent properties such as stiffness and yield stress of both materials. In this context, and following Darcy’s law, the bath materials solvent can diffuse into the ink material (pre-/post-crosslinking) under osmotic pressure causing the structure to swell. Simultaneously, this osmotic pressure can also result in the opposite, leading to ink shrinkage as components of the ink diffuse into the surrounding bath matrix.^[71,72]^ In both scenarios, these processes may contribute to an overall loss of structural fidelity and potential collapse of the construct. In addition, higher levels of dECM porosity were observed compared to materials bioprinted without a support bath (**Figure 3D**), which may contribute to the collapse of the constructs. However, additional studies would be necessary to validate the presented assumptions. Histological analysis of this cellularized material supported previous results showing a highly porous matrix composed of elastin and collagen, in which cells can be observed (**Figure 3E, S22**). We attempted to prevent cylinder collapse by increasing the thickness of the bioprinted dECM-based bioink, however this approach resulted in a loss of bioprinting fidelity (**Figure S21B**). We also evaluated increasing the UV exposure time for photo-crosslinking. Studies suggest that constructs bioprinted within support baths exhibit decreased stiffness compared to those bioprinted in air, attributed to partial crosslinking of the photo-polymerizable materials due to the bath’s incomplete transparency and inadequate UV penetration, thereby affecting the stiffness of the embedded pbiorinted constructs.^[73]^ However, based on the well-established fact that prolonged UV irradiation can compromise cell viability, we did not pursue this approach further.^[74,75]^ We finally characterized the growth of vSMCs over a 2-week period, observing concordant alpha-smooth muscle actin (α-SMA) staining and a network of vSMCs showing elongated extensions (**Figure 3F, G**).

**Figure 3.**
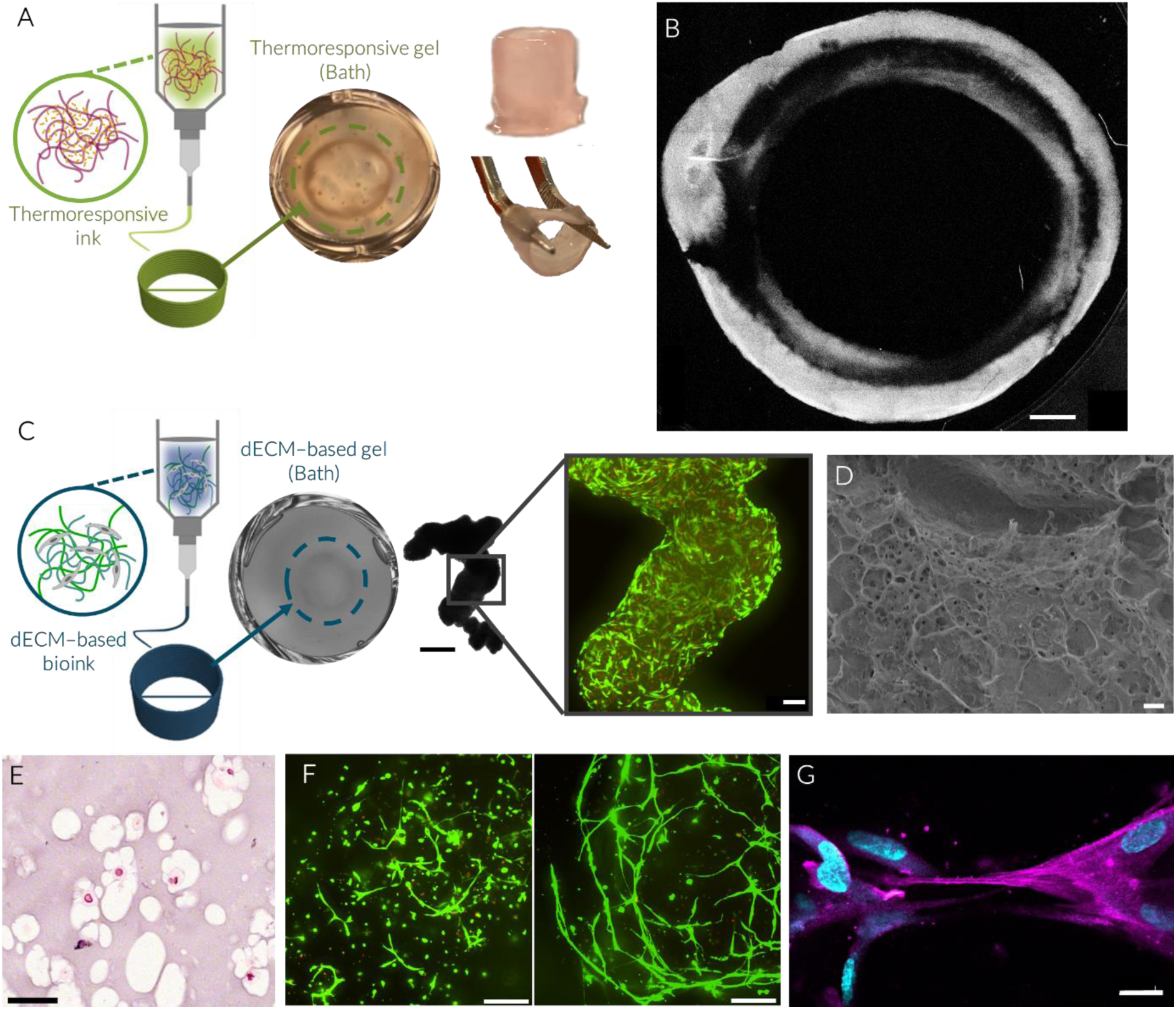
Embedded 3D bioprinting of single material hybrid ink and dECM-based bioink. A) Schematic of thermoresponsive ink printing and resulting 1 cm tall cylinders. B) Cross-section of the hybrid ink, sensitive to 2PEL thanks to embedded AuNRs (scale bar: 100 µm). C) Schematic of dECM-based bioink printing within the bath and resulting structure after 24 h in culture by brightfield microscopy (scale bar: 500 µm). hPASMC cell viability is observed using live/dead fluorescence confocal imaging, showing a MIP of 200 µm thick z-stack (scale bar: 100 µm). D) SEM images of dECM-based gel printing within the bath (scale bar: 50 µm). E) Histological evaluation of the embedded printed vSMC-based gels at day 7 including a representative image of H&E (scale bar: 50 µm). F) vSMC cell viability after, left to right, 24 h, and 14 days *in vitro*, showing extensive cell spreading of the cells within the dECM (scale bars: 200 µm). G) vSMCs cultured for 7 days within the construct showing α-SMA expression and nuclear staining (DAPI) (scale bar: 20 µm).

#### 2.2.2. Embedded 3D bioprinting of concentric multilayered cylinders

After confirming that both materials could be successfully printed and crosslinked within the support bath, we moved to bioprinting of multi-material concentric cylinders. To explore the minimal and maximal distances between the dECM-based and hybrid gel layers, thus ensuring their successful crosslinking without loss of bioprinting resolution, various concentric cylinder models of different diameters and thicknesses were designed (**Figure S23**). When the distance between the materials was not optimal, the layers failed to crosslink together, resulting in two separate cylinders (**Figure S23A**). Similarly, when the infill distance was greater than 0.5 mm, closed cylinders with no resolution were obtained (**Figure S23B**). Additionally, two printing approaches were tested: i) simultaneous, where a layer of each material was printed prior to changing the height of the deposited materials; and ii) sequential, where the entire cylinder of one material was bioprinted, followed by the entire cylinder of the other material. Within the sequential printing approach, we also studied variations in the order of bioprinting the model, whether starting with the inner or outer material first and vice versa. We observed that the highest bioprinting resolution was achieved using a simultaneous bioprinting approach in which the inner material was printed first, followed by the printing of the outer one (**Figure S24**). To achieve this, constant pressure for material extrusion was required and resulted in crucial for achieving high bioprinting fidelity. Indeed, both materials exhibit liquid behavior and similar rheological properties at RT, thus allowing us to fix the pressure at 20 kPa for both materials (**Figure S25**). Based on the adjustment of the model dimensions and the optimization of bioprinting parameters, we selected a 9 mm diameter cylinder for the outer layer and a 7 mm diameter cylinder for the inner layer, with an infill distance of 0.5 mm for the embedded bioprinting of the constructs with high reproducibility (**Figure S26**). The additional bioprinting parameters were those previously described and employed for the bioprinting of the individual materials, as summarized in **Table S2**.

Upon bioprinting with the optimized parameters, the multiple layers of the concentric cylinder were crosslinked together, resulting in self-standing and robust 3D bioprinted constructs (**Figure 4A, B**). Again, histological analysis confirmed the bilayered structure and clear biological differences in collagen and elastin content (**Figure 4C, S27**). Given that the dECM-based cylinder is bound to the hybrid layer, diffusion on the outer surface of the structure is restricted, which in turn limits osmosis-controlled swelling and shrinking mechanisms. As hypothesized, upon crosslinking with the hybrid layer, the cell-containing dECM-based bioink maintained its cylindrical form and remained securely bound to the external layer for over 7 days. SEM imaging confirmed uniformity in the contact between the layers, with high magnification SEM suggesting polymeric fiber interactions due to the crosslinking procedure (**Figure 4D, Figure S28**). Via 2PEL and fluorescence confocal imaging highlighting the AuNP-containing hybrid inks and actin-stained cell-containing dECM-based bioinks, respectively, we were able to observe the bi-layered model with high resolution (**Figure 4E**). vSMCs were homogeneously distributed throughout the cylinder, displaying high cell viability up to 14 days *in vitro* (**Figure 4F,G**). Results from vSMC metabolic activity testing in single and multi-layered cylinders suggested that single cylinders better supported cellular activity, potentially due to the change in dECM-based bioink stiffness due to its chemical cross-linking with the outer hybrid gel layer (**Figure S29**).

**Figure 4.**
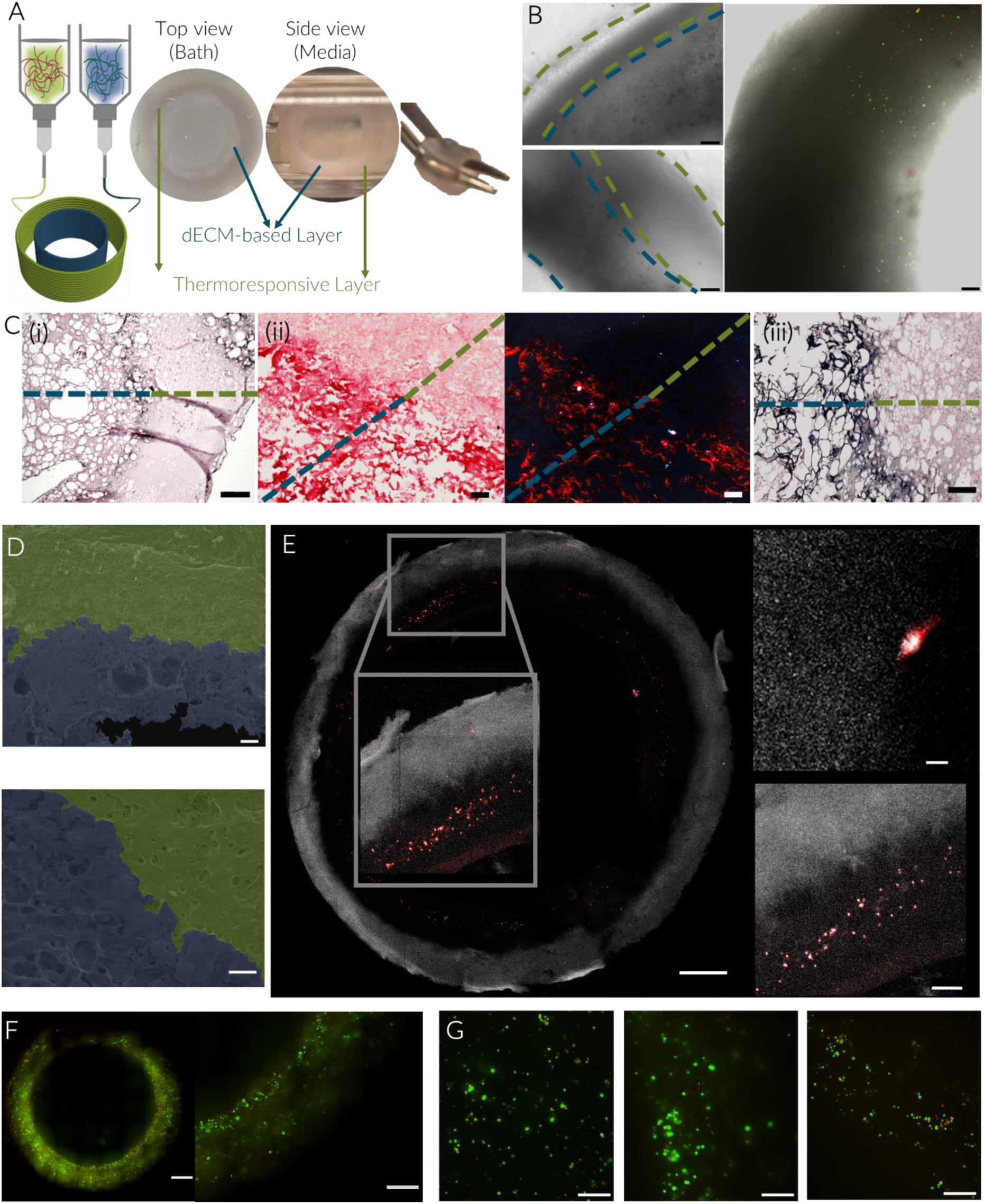
Multi-material embedded 3D bioprinting of concentric cylinders. A) Schematic representation of the 3D printed model and images of the printed structures i) within the bath, ii) after its removal, iii) incubation with fresh media, and iv) out of the printing wells. B) Brightfield microscopy showing layers and live/dead stained vSMCs (scale bars: 200 µm). C) Histological analysis of multilayered constructs including representative images of (i) H&E, and analysis of the structure and composition through the study of the presence of (i) collagenous fibers by brightfield images (left) and polarized light (right) of Picrosirius Red staining and (iii) Verhoeff Van Giesson stain showing elastic fibers (scale bars: (i); 200 µm, (ii); 50 µm, and (iii); 100 µm). D) Structural characterization of the printed constructs via SEM, including the hybrid gel and dECM-based layers highlighted in green and blue, respectively. (scale bars: top, 200 µm; bottom, 50 µm). E) Combined 2PEL and fluorescence confocal microscopy showing the hybrid gel layer (grayscale) and actin-labeled vSMCs (red) (scale bars: left, 1000 µm; top right, 20 µm; bottom right, 200 µm). F-G) Cell viability, assessed via live/dead cell fluorescence imaging, 24 h post-printing (F) (scale bars: 1000 µm and 500 µm), and subsequently at day of 1, 7, and 14 *in vitro* (G) (scale bars: 200 µm). All confocal microscopy images are Maximum Intensity Projections (MIPs) of 200 µm thick z-stack.

## 3. Conclusions

We have demonstrated the potential of CLADDING embedded bioprinting for the fabrication of a cellularized concentric multimaterial layered artery model. Two novel bioinks, compatible with embedded bioprinting followed by successful photocrosslinking within the bath, were generated for the development of the *tunica adventitia* and *tunica media* layers of blood vessels. Specifically, for the outermost *tunica adventitia* layer of the model, a thermoresponsive elastic hybrid ink has been developed, which can impart stimuli-responsiveness to the model and structural support to the adjacent cell-containing inner layer. For the *tunica media* layer, a dECM-based bioink, derived from porcine arteries and combined with vSMCs, which presented high cell viability and cell spreading over time has been described. With the optimized ink formulations and refined printing and crosslinking parameters, the generation of self-standing constructs with simultaneously crosslinked concentric cylinders, supportive of vSMC growth, has been achieved. These novel constructs open new possibilities in *in vitro* model hybrid designs. Based on our previous work in which we observed changes in the mechanoadaptative signature when subjecting the constructs to temperatures above and below the LCST, modifying the material stiffness, a similar approach could be implemented here. To understand cellular responses to mechanical forces and matrix stiffness, further analysis is needed, especially considering that the constructs were cultured at 37°C - a temperature at which the hybrid gel remains contracted, potentially exerting additional pressure and compactness on the cells.

Additionally, it provides the possibility in future work to implement cyclic expansion and contraction of the multilayered model by NIR laser stimulation to modulate the stiffness of the material through plasmonic heating of the thermoresponsive layer, replicating the pulsation of native arteries as previously reported.^[15]^ Overall, the versatility of both the hybrid ink and dECM-based bioink, in combination with embedded bioprinting, can be exploited to replicate a wide range of native structures and their corresponding functions, opening new avenues in tissue engineering.

## 4. Experimental Section

### Materials

Hexadecyltrimethylammonium bromide (CTAB, ≥99.0%), hydrogen tetrachloroaurate trihydrate (HAuCl_4_·3H2O, ≥99.9%), silver nitrate (AgNO_3_, ≥99.9%), L-ascorbic acid (AA, ≥99%), sodium borohydride (NaBH_4_, 99%), N-isopropylacrylamide (NIPAm, 97%), poly(ethylene glycol) diacrylate (PEGDA, average Mn=700), Poly(ethylene glycol) dimethacrylate (average Mn=550), Lithium phenyl-2,4,6-trimethylbenzoylphosphinate (LAP, ≥95%), gelatin from porcine skin (gel strength 300, type A), agarose (wide range, for molecular biology), gum arabic from acacia tree, Pluronic® F-127, methacrylic anhydride (MA, ≥94%) dialysis tubing cellulose membranes, Tris(2,2’-bipyridyl)dichlororuthenium(II) hexahydrate, sodium persulfate, DPX mountant for histology, papain from papaya latex (lyophilized powder, ≥10 units/mg protein), sodium deoxycholate (SDC, ≥95%), sodium dodecyl sulfate (ReagentPlus®, ≥98.5%), DNase I (native from bovine pancreas), pepsin (pepsin A from porcine gastric mucosa), acetic acid (glacial, ACS reagent, ≥99.7%), peracetic acid (PAA, about 38-40%), ethanol (EtOH), sodium chloride (BioXtra, ≥99.5%), TWEEN® 20, Triton™ X-100, bovine serum albumin (BSA), calcium chloride and D-Cysteine hydrochloride monohydrate (≥98%), picric acid solution, Direct Red 80, Eosin Y Solution (alcoholic, with Phloxine), Elastic Stain, sodium hydroxide (NaOH) and hydrochloric acid (HCl), were all purchased from Merck. Ethanol (EtOH, absolute) for support bath preparation was purchased from Scharlau. Gelatin type B from bovine bone (type II) 250 LBB8 was obtained from Rousselot Biomedical. 0.5% trypsin/EDTA solution (10X), penicillin-streptomycin (P/S, 10,000 U/mL), 4′,6-diamidino-2-phenylindole (DAPI, 1 mg/mL), actin-555 ReadyProbes, ethidium homodimer-1 (EthD-1), calcein acetoxymethyl (Calcein AM), PrestoBlue^TM^ cell viability reagent, alamarBlue^TM^ cell viability reagent, paraformaldehyde (PFA), lactate dehydrogenase (LDH, CyQuant assay) kit, Quant-iT™ PicoGreen™ dsDNA Reagent, deoxyribonucleic acid from calf thymus and ethylenediaminetetraacetic acid (EDTA, 0.5 M PH 8.0) and Alpha-Smooth Muscle Actin Antibody (MA1-06110), were all purchased from Thermo Fisher Scientific. Donkey F(ab’)2 Anti-Mouse IgG H&L (Alexa Fluor® 647, ab150103), was purchaser from abcam. CellWax Plus was purchased from Cellpath. Harri’s Hematoximin and xylene were purchased from Labolan. Human Pulmonary Artery Smooth Muscle Cells (vSMCs, ATCC-PCS-100-023), Vascular Cell Basal Media (ATCC-PCS-100-030), and Vascular Smooth Muscle Cell growth supplements (ATCC-PCS-100-023) were purchased from ATCC. All chemicals were used as received and Milli-Q water was used in all experiments. All glassware used for AuNRs synthesis was washed with aqua regia, rinsed with water and dried before use.

## Hybrid ink preparation and characterization

### Synthesis of GelMA

Gelatin was functionalized with methacrylate groups for the synthesis of gelatin methacryloyl (GelMA). 5 g of gelatin (type A) were dissolved in 100 mL of PBS (pH = 7.4) upon heating to 50 °C and under vigorous stirring. Once the gelatin was completely dissolved, methacrylic anhydride (MA) (0.8 mL/g gelatin) was added and the mixture was stirred at 500 rpm for 3 h at 50 °C. After the reaction was completed, the solution was dialyzed using cellulose membranes (MW cut-off 3.5 kDa) against distilled water at 45 °C for 5 days to remove the unreacted reagent. Methacrylated gelatin (GelMA) was recovered after freeze-drying for 72 h. The degree of methacrylation (∼80%) was estimated by H-NMR by comparing the integration of the signals of the methyl groups around 1.95 ppm with the ones corresponding to the methylene group at 5.6 and 5.35 ppm (**Figure S30**). Lyophilized GelMA was dissolved in PBS at a final concentration of 10 w/v % and stored at 4 °C until combined with the other components for the formation of the bioink.

### Nanoparticle Synthesis and characterization

Nanoparticle synthesis was conducted using a microfluidic flow reactor setup, following a previously established protocol. ^[76]^ Briefly, seeds were prepared by the rapid reduction of a 20 mL HAuCl_4_ 1 mM solution in 100 mM CTAB, using a 2.4 mL NaBH_4_ 12 mM aqueous solution. Two HPLC pumps and a total flow rate of 11.2 mL/min were maintained within the reactor for the preparation of the seeds, which were kept at RT for 30 minutes to remove the borohydride excess. All three pumps of the reactor were employed for the synthesis of AuNRs. For the growth of the AuNRs, a 50 mL solution of 100 mM CTAB containing 1 mM HAuCl_4_ and 0.24 mM AgNO_3_ under acidic conditions (pH 3 ∼ 4) was mixed with a 25 mL solution of 3.2 mM ascorbic acid in 100 mM CTAB. Following Au(III) pre-reduction to Au(I) in the mixing tube, 25 mL of 100 mM CTAB containing 800 μL of seeds were added, with a 20 mL/min total system flow rate. The resulting solution was left undisturbed for 4 h before centrifuging the colloidal dispersion at 8000 rpm for 20 minutes and redispersing the AuNRs in a 1 mM CTAB solution. This method resulted in AuNRs with a maximum absorbance at 790 nm, observed in the UV−vis−NIR extinction spectra (Agilent 8453 UV−vis diode-array spectrophotometer). AuNR characterization via Transmission electron microscope (JEOL JEM-1400PLUS) operating at 120 kV, was conducted by drop casting on a carbon-coated 400 square mesh Cu grid. The TEM images revealed that the AuNRs exhibited high monodispersity, with an aspect ratio of 3.4 and final dimensions of 82.8 ± 7.1 nm by 24.4 ± 3.6 nm (**Figure S1**).

### Polymeric ink development

The final hybrid ink was prepared by mixing, in a 3:1 ratio, two concentrated solutions including a stock of NIPAm, PEGDA and PEGDMA and the other one of GelMA with AuNRs. Different inks with varying concentrations of PEGDA and PEGDMA were designed, which were then integrated with the AuNR containing GelMA matrix. For reaching the desired final concentration, 21.33 % (w/v) NIPAm and 8 w/v % PEGDA were added into PBS (10 mM, pH 7.4), and dissolved upon magnetic stirring and sonication at RT. 2.66 % (v/v) PEGDMA was then introduced to the first solution, and complete homogenization was achieved upon centrifugation at 5000 rpm for 5 minutes. In control gels without PEGDMA, 10.66 % (w/v) PEGDA was added to the NIPAm, maintaining in both cases, the combined sum of both PEG derivatives in 10.66 % (w/v). The resulting formulations were stored at 4 °C until needed. For the GelMA-AuNRs solution, 20 % (w/v) GelMA was dissolved in PBS under vigorous stirring at 45 °C. The AuNR solution was gently washed twice by centrifugation at 8000 rpm for 10 minutes and resuspended in milliQ water to eliminate the excess of CTAB. The NP concentration was adjusted during washing to achieve the desired concentration of 2 mM Au^0^ when combined with the GelMA. The washed AuNR solution was added drop by drop to the dissolved GelMA gel under stirring, ensuring that the color of the final gel matched the color of the AuNRs in solution. For the final gel preparation, the GelMA - AuNR solution was added drop by drop to the concentrated hybrid ink mixture under continuous stirring in a 3:1 ratio. The resulting formulation, containing 16 w/v % NIPAm, 8-6 % (w/v) PEGDA, 0-2 % (v/v) PEGDMA, 5 % (w/v) GelMA and 0.5 mM AuNRs, was stored at 4 °C until needed. Immediately prior to use, the photoinitiator LAP was added to the gel at a final concentration of 0.1 % (w/v).

### Mechanical characterization

An Instron universal testing system (68SC-2 Single Column Table Model) was employed for the mechanical characterization of the hybrid ink. Compression and tensile stress-strain tests were performed using discs of 3 mm in diameter and 1 mm thickness, and dog-bone shaped samples of 15 mm length, 2 mm width and 1 mm thickness. In both tests, an incremental strain was applied in opposite directions at a constant rate of 0.5 mm/min. The young’s modulus was calculated by the linear regression of the tensile test stress-strain curve. All measurements were done in triplicate.

### Differential scanning calorimetry (DSC)

To characterize the transition temperature of the ink and verify that the LCST of the formulation is close to the physiological body temperature, DSC was performed. A dynamic sweep from 10 - 50 °C at 5 °C/min in a nitrogen atmosphere was conducted where the LCST of the final gel was determined by the average inflection points of the temperature change profiles of the DSC curves. The measurements were done in triplicate. *Thermal characterization.* To homogenously irradiate the AuNR-hydrogels, a SuperK Extreme white light ultra-broadband supercontinuum class four laser source (NKT photonics) delivered in a single mode fiber was employed. A 78 MHz seed laser with light frequencies comprising the range from 400 to 2400 was emitted in a single spatially coherent beam, that was coupled to a complete fiber delivery system (FD7-PM) paired with a SuperK CONNECT beam alignment device. The system was connected to the module SuperK VARIA, which provides a filtered output channel using a variable bandpass filter. The bandpass filter was adjusted within the range of 730 – 850 nm, specifically chosen for its higher power output. The distance from the laser fiber output to the sample was adjusted with a laser aligning card to ensure that the spot size measured 1 cm^2^. The intensity of the laser was set to 100% in all cases to maintain uniformity across experiments. A pulse generator was employed to regulate the pulsation of the supercontinuum laser, which was configured in external current mode. A pulsed waveform with an amplitude set at its maximum of 20 Vpp and a frequency of 10 Hz, featuring duty cycles of 15%, 17.5%, and 20%—each representing the percentage of one period during which the signal is active—was chosen for the cyclic heating of the material. Hydrogel discs, measuring 1 cm in diameter and matching the laser spot size, were immersed in PBS in 24-well plates to simulate cell culture conditions and placed on a hot plate set to 35 °C. To monitor the temperature changes, the system was connected to a thermal camera (FLIR E8 pro), allowing the *in situ* recording of the temperature during heating. Finally, to achieve rapid and cyclic heating of the hydrogels around the LCST temperature range, the amplitude was set to its maximum of 20 Vpp, operating at a frequency of 10 Hz with a heat plate maintaining a baseline temperature of 35 °C, and a 20 % duty cycle allowing the temperatures to stay within the LCST range.

## dECM-based bioink development and characterization

### ECM decellularization and bioink formulation

The main pulmonary artery and smaller arterial capillaries were extracted from whole porcine lungs obtained from the local animal facility, treated with a 0.5 % (w/v) sodium dodecyl sulfate (SDS) and 0.5 % (w/v) sodium deoxycholate (SDC) solution for 72 h with intermediate 1 h washes in 0.9 % (w/v) NaCl to remove the residual detergent. The decellularization treatment was followed by a Deoxyribonuclease I (DNase I) treatment (40 U/mL) for an additional 24 h at 37 °C. Following the decellularization, tissues were sterilized using a 0.1 % (v/v) peracetic acid (PAA) solution in 4 % (v/v) ethanol (EtOH) for 2 h, followed by two 15-minute washes in PBS and two additional 15-minute washes with deionized water. Subsequently, the sterilized decellularized tissues were lyophilized, pulverized and the final dECM solution was obtained by digesting 10 mg of dECM powder per mL of 0.5 M acetic acid containing 2 mg of pepsin/mg of dECM at RT for 72 h. The pH of the solution was neutralized to a physiological pH 7.4 on ice to stop the digestion of pepsin. Finally, the resulting solution was stored at 4 °C to avoid its gelation until its use.

### Histology of the ECM and dECM

The ECM and dECM tissues were fixed in 4 % (v/v) PFA overnight at 4 °C, washed in PBS, dehydrated through graded washes of ethanol in water (50, 75 and 100%), washed twice with xylene and finally embedded in liquid paraffin at 60 °C for 1 h for subsequent staining. Frozen blocks were stored at - 80 °C until the histological sections were made with an HM 355S microtome. After de-paraffination, rehydration and permeabilization of the tissues with 0.3 % (v/v) Triton X-100 for 30 minutes, the 6 μm tissue sections were stained with 1 μg/mL DAPI for 30 minutes and mounted using Fluoromount-G. In addition, the tissue sections were subjected to Hematoxylin/Eosin (H/E), Picrosirius Red and Venhoerf Van Giesson staining with methods followed and adapted from described procedures and mounted in DPX.^[77]^

### Liquid chromatography-mass spectrometry (LC-MS) proteomics

Proteomic analysis was performed by the Proteomics Platform service at CIC bioGUNE. Briefly, 250 µL of extraction buffer (7 M Urea, 2 M Thiourea, 0.4% 3-[(3-cholamidopropyl) dimethylammonio]-1-propanesulfonate (CHAPS), 200 mM DTT) was added to each sample. Mechanical homogenization was performed in FasyPrep-24^TM^ 5G (MP Biomedicals). Samples were sonicated and centrifuged (13.000 rpm for 15 minutes) and supernatants were transferred to another tube. 50 µL of sample corresponding to native tissue and 250 µL of bioink were transferred to other tubes for overnight acetone precipitation at -20 C. Samples were centrifuged (13.000 rpm for 15 minutes), supernatants were discarded, and pellets were resuspended in 100 µL of extraction buffer. Protein content was quantified following the Bradford method. 50 µg of protein from each sample was trypsin (GOLD, Promega) digested following FASP method with minor modifications.^[78]^ The resulting peptides were speed vacuumed, resuspended in 0.1% trifluoroacetic and desalted and concentrated by using reverse phase microcolumns (C18 OMIX, Agilent). 200 ng from each sample was loaded on EVOSEP ONE (Evosep) coupled on-line to PASEF powered timsTOF Pro (Bruker) tandem mass spectrometer. Data Dependent Acquisition was applied with a 30 SPD method. The mass spectrum raw datasets were analyzed using the FragPipe (v22.0) free software for protein identification and quantification.^[79]^ Then, to identify differential proteins between conditions, Perseus software^[80]^ (https://www.maxquant.org/perseus/) was employed to carry out label free comparative analysis, considering proteins identified with at least one unique peptide at FDR<1% (PSM-level).

Intracellular and membrane proteins were discarded, and only ECM-related proteins were plotted and analyzed according to MatrisomeDB.^[81–83]^ Venn diagrams were constructed using Flaski, a collection of web apps for data analysis and visualization.^[84]^ Gene ontologies (GO) and Reactomes were obtained with g:Profiler, an interactive and collaborative web-based tool used for functional enrichment analysis that integrates various biological databases. Fisher’s one-tailed test, also referred to as cumulative hypergeometric probability, is used in this web-tool to calculate the p-value, which measures the randomness of the overlap between the query and the ontology term. This p-value indicates the probability of the observed intersection, as well as the probabilities of all larger, more extreme intersections. Principal component analysis (PCA) was performed with Metaboanalyst 6.0 software.^[85]^ The statistical significances of the group patterns are evaluated using PERMANOVA. The distributions were computed using the Euclidean distance based on the PCs. Biological interpretation of the results was conducted using GeneCards^[86]^.

### Image acquisition

The stained histological sections were imaged with Axio Observer microscope (Zeiss). Fluorescent samples were imaged with a Colibri LED module using 365 nm excitation for DAPI. H/E, Picrosirius Red and Verhoeff Van Giesson-stained sections were acquired using a brightfield and 305 Axiocam camera. Collagen fibers stained with Picrosirius Red were imaged through polarized light. The EC Plan Neofluar 10x (numerical aperture NA, 0.30, working distance WD, 5.3 mm) objective was employed for image acquisition which was controlled by the Zen Blue software (v.2.3).

### Collagen quantification

Optical anisotropy or birefringence phenomena arises upon the interaction of polarized light with anisotropic materials such as collagen. The triple helix structure of collagen causes the light waves to divide into two perpendicular components, which will travel at a different speed through the material, resulting in birefringence. This effect enables the visualization of collagen’s structure, alignment, and density within tissues. To determine the percentage of collagen in the tissue and compare the native and decellularized tissue samples, the brightfield image (showing the entire tissue) and polarized light images (highlighting only the collagen) of the Picrosirius red staining were analyzed using ImageJ. Briefly, brightfield images were converted to black and white for optimal visualization and a threshold was adjusted to cover all tissue areas while excluding non-tissue elements. Particle analysis was then performed, with the size set from 200 to infinity, and the results are manually refined to remove any irrelevant particles. The total area of the tissue was calculated as the sum of all identified particles. This process was repeated for the polarized image and the collagen content was determined using the formula:

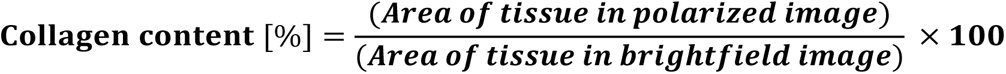

This calculation was carried out for both native and decellularized tissues to assess collagen retention post-decellularization.

### DNA quantification

Genomic DNA was extracted from the ECM and dECM tissues by digesting 5 mg of each tissue in 500 μL of digestion buffer (6 mM cysteine HCl, 6mM EDTA and 4 U/mL of papain in PBS) for 72 h at 37 °C. DNA was manually precipitated with ethanol as described in Green et al. ^[87]^ Quant-iT™ PicoGreen™ dsDNA Reagent was employed for the quantification of double-stranded DNA through fluorescence, following the manufactures instructions and including calf thymus DNA in 1× Tris-EDTA buffer as a standard. Samples in 96-well plates were excited at 492 nm and emission was recorded at 535 nm. The data has been averaged by 7 biological samples including 3 technical replicas of each (N = 7, n = 3) and expressed as the mean values and standard deviation of ng of DNA per mg of tissue.

### Biocompatibility of dECM gels

The cytotoxicity and biocompatibility of the dECM was assessed by a lactate dehydrogenase (LDH) assay kit complemented with an AlamarBlue assay. hPASMC were seeded at a density of 31,250 cells/cm^2^ and cultured with 100 µL of fresh media on 96-well plates. The crosslinked artery dECM derived gels of different concentrations were incubated in complete vascular cell media at 37 °C for 24 hours. The cell media of the cells in the plate were then replaced with conditioned medium and incubated for additional 24 h. Lysis buffer and fresh media were added as the positive and negative controls, respectively. For the LDH Assay, 50 µL of substrate mix was added on top of 50 µL of cell media in an optically clear 96-well plate, followed by a 30-minute incubation at RT. Subsequently, 50 µL of stop buffer was added, and absorbance readings were taken at 490 nm and 680 nm. For the AlamarBlue Assay, cells cultured with the conditioned media for 24h, were incubated in AlamarBlue containing basal vascular cell media (1/10) for 3h at 37 °C. The level of fluorescence read at 570 nm (excitation) and 585 nm (emission) correlates directly with the quantity of viable cells, reflecting their metabolic activity. Data shown in graphs were the summary of three biological replicates, with three technical replicates and analyzed using GraphPad Prism 8 software (GraphPad, San Diego, CA).

### Cell culture

Human Pulmonary Artery Smooth Muscle Cells (pPASMCs) were cultured in Vascular Cell Basal Media supplemented with vascular smooth muscle growth supplements and used between passages 3 and 12. The cells were trypsinized using 0.5x trypsin-EDTA. *dECM – GelMA bioink development.* Different bioink compositions were prepared by varying the concentration of the dECM, GelMA and LAP for cell viability and structural fidelity studies. The GelMA concentration range was extended from 5 to 2.5 % (w/v) and prepared by diluting the 10 % (w/v) stock in PBS. Working on ice, 0.1 %, 0.25 % and 0.5 % (w/v) dECM gels were prepared from 1 % (w/v) dECM solutions by diluting the matrix in PBS. The photoinitiator, LAP was freshly added to the formulation from a 1 % (w/v) stock prepared in PBS, reaching a final concentration of 0.1 w/v % in the bioink. To ensure sterility and avoid contamination the bioink was filtered with a 0.2 µm pore filter. hPASMC were mixed with the dECM-based bioink at a concentration of 1.5 × 10^6^ cells/mL to form the vSMC bioink.

### Rheology

Rheological properties of the thermoresponsive and the cell-free dECM-based inks and gels were assessed using a MCR 302 rheometer (Anton Paar) equipped with a 25 mm diameter parallel plate geometry and a solvent trap to prevent water evaporation. The gap for measurements was adjusted to the type of measurement and all tests were carried out in triplicate. Flow curves were performed to characterize the shear stress and viscosity for shear rates from 0.1 to 1000 s^-1^ at the printing temperature, 20 °C and with a gap of 0.25 mm. To simulate the crosslinking of the different materials upon bioprinting, initially the inks were left at rest for 120 s, allowing for observation of its baseline properties, corresponding to the material lying in the syringe barrel prior to bioprinting. The samples were then irradiated with UV light (OmniCure Series 1500, Excelitas Technologies) for 30 s @17.25 mW/cm^2^ power density and left at rest for additional 400 s, representing the time the gels were in the bioprinting bath before their incubation at 37 °C. For the dECM-based bioink, the samples were subsequently subjected to 37 °C for 400 s, for assessing the thermal crosslinking of the material. To gain insights into the properties of the thermoresponsive gel, crosslinked samples were subjected to amplitude sweeps from 0.001 to 100 %, at a constant angular frequency of 1 Hz at 37 °C, to determine the linear viscoelastic region of the material employing a gap of 1 mm. To determine the storage and loss moduli of the different samples, frequency sweeps were carried out from 500 to 0.1 rad/s, at a fixed strain of 0.1 %, included the viscoelastic region. Temperature ramps were performed from 20 to 50 °C at 1 °C/min, 1 Hz and 0.1 % strain.

## Elemental and microscopical characterization

Scanning Electron Microscopy (SEM) of the thermoresponsive gel and dECM-based gel was performed to determine the microstructure of the bioprinted constructs using a Field Emission Scanning Electron Microscope – FESEM (JEOL JSM-IT800) operating at an accelerated voltage of 2 kV and a working distance of around 4 mm. The SEM images were falsely colored using Inkscape 1.2 software.

## Bioprinting of models

A multiheaded 3D Discovery bioprinter (RegenHU, Switzerland) was used to bioprint the structures without a supporting bath and within the gelatin and agarose baths. The G-codes for bioprinting the structures, including 7 and 5 mm in diameter and 1 cm in height cylinders, were produced using BIOCAD software. The stimuli-responsive material was bioprinted through a precision plunger dispenser at a constant volume flow rate of 2 μL/s using a conical plastic needle with an inner diameter of 0.2 mm (Nordson). The material was bioprinted at RT or at 5 °C, and a collector plate speed of 5 mm/s was employed for the bioprinting. The dECM–based bioinks were extruded through a dispensing plastic needle with an internal diameter of 0.2 mm at RT, pressures of 0.045 MPa, and printing speed of 5 mm/s. A light-curing printhead, operating at 365 nm with a fixed output power of 500 mW, was used to cure the constructs. The irradiation time and frequency were optimized depending on the bioprinting approach being conventional or gelatin or agarose support bath assisted. Overall, crosslinking times ranging from 30 seconds to a maximum of 7 minutes were employed and depending on the case, the irradiation was applied every layer, every certain number of layers or upon the completion of the bioprinting of the construct.

## Preparation of support baths

### Gelatin slurry

A well-established protocol was followed for the preparation of the gelatin slurry support bath.^[88]^ Briefly, 4.5 % (w/v) gelatin (Type A) was dissolved in 150 mL PBS at 60 °C under stirring in a 500 mL mason jar. The solution was covered and incubated at 4 °C for 12 h. Subsequently, 350 mL of PBS at 4 °C was added, and the mixture was blended 120 s in a consumer-grade blender. The resulting blended gelatin slurry was transferred into 50 mL Falcon tubes and centrifuged at 4200 rpm for 2 minutes, causing the slurry particles to settle. The supernatant was removed, replaced with PBS at 4 °C and vortexed to re-suspend again. This centrifugation and washing cycles were repeated until no bubbles were visible at the top of the supernatant, indicating the removal of most soluble gelatin. At this point, the solution was stored at 4 °C until needed or directly employed for embedded printing.

### Agarose slurry

A similar protocol was followed for the preparation of an agarose slurry support medium.^[26]^ Agarose (3.33 % (w/v)) was dissolved in 150 mL of PBS at 60 °C under stirring. After storing the dissolved agarose solution overnight at 4 °C, an additional 350 mL of PBS were added to the mixture, which was placed at –20 °C for 20 minutes. The gel was subsequently blended for 120 s in a consumer-grade blender and transferred to 50 mL Falcon tubes for the washing cycles. The same centrifugation conditions as for the gelatin slurry preparations were employed and repeated until a refined slurry was obtained. Again, the agarose slurry gel was stored at 4 °C or directly used at this point.

*The support bath for CLADDING bioprinting* was prepared through a coacervation process, following an established protocol.^[32]^ The coacervate preparation involved mixing gelatin type B from bovine bone and gum Arabic (4:1) in deionized water, which constituted 50 % of the total solvent volume. The mixture was then heated to 45 °C and vigorously stirred until complete dissolution, followed by the addition of the remaining 50 % of the solvent, consisting of absolute ethanol and 2.5 % (w/v) Pluronic F-127. The pH was then adjusted to 5.8, and the solution was stirred for 15 minutes before being left undisturbed at RT for 6 h. This led to the precipitation of gelatin-arabic gum microparticles. The micro-coacervation process was completed by placing the solution at 4 °C overnight, and removing the liquid equilibrium phase, located above the coacervate, formed at the bottom of the flask. With special care to not disturb the coacervate, the gel was transferred to 50 mL Falcon tubes and stored at 4 °C before use. For optimal bioprinting, the hydrophobicity of the supporting bath was enhanced through a 3-step washing process including two centrifugations at 1000 g for 5 minutes and a final one at 2000 g. For effectively compacting the microparticles to yield an optimized supporting bath, PBS (20 mM), CaCl_2_ (20 mM) and Tween20 (5 % (v/v)) were tested as washing solutions.

### Absorbance of support baths

A CLARIOstar plate reader (BMG Labtech) was employed to measure the absorbance in the range of 300 – 500 nm of the different support baths at RT. In addition, for the finally selected embedding bath for CLADDING bioprinting, the overtime absorbance of the material was measured at 37 °C. In both cases, to replicate the experimental conditions, 2 mL of washed suspension bath solutions were added into 24-well plates.

### CLADDING bioprinting

The bioprinting setup included two N_2_ pressure-based systems for the simultaneous deposition of the materials controlled by the bioprinter (BioScaffolder 3.1, GeSiM, Germany), a UV light source, and a 24-well plate loaded with 2 mL of support bath, employed as the bioprinting platform (**Figure S31**). The environment, bioprinting support and cartridge temperatures were not controlled, but the temperature of the room was maintained at ∼ 20 °C. Single and concentric cylinder models of various dimensions were designed using 3D Builder software. Based on these 3D models, multiple G-codes were developed to improve the fidelity, stability and reproducibility of both single and multilayered self-standing cylinders. Both inks were loaded into 3 mL syringe barrels and printed with 2.54 cm long metallic needles of 2-3 mm diameter (Nordson EFD, Inc) at a pressure of 20 kPa. Single-layered thermoresponsive cylinders of 9 mm in diameter and 1 cm in height and dECM-based cylinders, containing one and two circles per layer, of 5-7 mm in diameter and of 0.5 cm in height and, were bioprinted at collector printing speeds of 15 and 10 mm/s, respectively. For both materials, a strand width that matches the nozzle diameter, a strand height of 0.3 mm, and a 60 °C angle change between the layers were implemented across the layers to prevent gap formation, which could compromise the integrity of tall, self-standing cylinders due to insufficient material deposition at the start of each layer. For the multimaterial bioprinting of concentric cylinders, 0.25 mm diameter nozzles were employed. Models combining an outer layer of 9 mm diameter with inner layer diameters of 5, 6, and 7 mm, and bioprinted with infill distances ranging from 0.5 mm to 2 mm, were evaluated. A light-curing printhead operating at 365 nm with an output power of 11.6 mW/cm² was employed to crosslink the inks post-printing. The curing process involved simultaneous crosslinking of the layers by exposing the entire printed construct to 365 nm light for one minute after printing was completed. This exposure was divided into two phases: 30 seconds from the top and 30 seconds from below ensuring a thorough and uniform curing of the printed model.

The bioprinted constructs were subsequently incubated at 37 °C for 45 minutes, enabling the thermal crosslinking of the dECM component and melting of the support bath. It was noted that the bath transitioned from being opaque at RT to fully transparent when incubated at 37 °C, leading to the material melting (**Figure S15**). By monitoring the absorbance of the material over time, we noted a decrease in the bath’s absorbance beginning at 25 minutes, stabilizing near 0 approximately 40 minutes after incubation at 37 °C. Based on these findings, we chose to start washing the bath and supplying the tissue analogues with media after 45 minutes of incubation at 37 °C, ensuring the thermal crosslinking of the dECM-component and melting of the support, maximizing cell viability. At this point the bath was removed, the bioprinted constructs are washed with PBS and warm fresh cell media was added for cell culture.

## Cell growth and characterization in 3D model

The multi-layered bioprinted constructs were incubated at 37 °C and 5 % CO_2_ in a humidified incubator, replacing cell media every 2-3 days. A stereoscope was used to observe changes in cell morphology over time. Cell survival within the bioprinted single dECM-based cylinder and the multilayered concentric layer model was evaluated with live/dead staining assay combined with PrestoBlue® assay. 3D models were incubated with a solution consisting of 2.5 μM of ethidium homodimer-1 and 1 μM of calcein for 1 h, washed with complete fresh media and imaged with a fluorescence microscope (Eclipse Ti-E Nikon, Japan). For the PrestoBlue® cell viability assay, the 3D models were incubated with the PrestoBlue® solution containing basal vascular cell media (1 in 5 dilution) for 2 h at 37 °C. The supernatant was collected and transferred to an optically clear 96-well plate. The level of fluorescence read at 560 nm (excitation) and 590 nm (emission) correlates directly with the quantity of viable cells, reflecting their metabolic activity. After the measurement, the 3D model was washed with PBS until no traces of the PrestoBlue assay solution were observed and the constructs were cultured in fresh media to study their evolution in time. To achieve higher resolution and differentiate better the morphology of the cells embedded in the constructs, fluorescence confocal imaging of the samples was conducted. For nuclear and actin staining, samples were fixed (using 4 % (w/v) PFA) at various time points and stained using DAPI (1/500) and Actin 555 ReadyProbe (1/60) at 4 °C overnight. For the immunofluorescence assays, samples were fixed (using 4 w/v% PFA) and permeabilized with 0.25% Triton X-100 in PBS for 1 h at RT. Blocking was performed using a solution of 1 % (w/v) bovine serum albumin (BSA) and 0.1 % (v/v) Tween-20 in PBS (PBST) for 1 h at RT, ensuring maintenance of humidity preventing the drying of the models. The primary antibody, staining against alpha smooth muscle actin (MA1-06110) was diluted in blocking solution obtaining a final concentration of 10 µg/mL and incubated overnight at 4 °C. After washing in blocking solution, the samples were incubated with Alexa647-conjugated secondary antibody (ThermoFisher) overnight at 4 °C while protected from light at a final concentration of 10 µg/mL. The samples were washed with PBS, DAPI (1/500) was added for nuclear staining at a concentration of 1:500 in milliQ water, and samples are washed with PBS prior to imaging. All washes were performed for at least 30 minutes each to ensure thorough removal of excess reagents. Confocal fluorescence and multiphoton imaging were conducted using a Zeiss LSM 880 microscope equipped with argon, DPSS, HeNE and MaiTai multiphoton lasers. Images were processed with ZEN and ImageJ software.

### Statistical Analysis

Bar graphs and point graphs display mean value ± SD. Linear regression analysis was performed to determine the Young’s Modulus of the materials, with statistical significance levels at p < 0.0001. All analyses and graphs were performed using GraphPad Prism 8 software (GraphPad, San Diego, CA).

## Supporting Information

Supporting Information is available.

## Acknowledgements

UAL thanks the Basque Government’s Predoctoral Programme (PRE_2023_2_0014). MCIN/AEI/10.13039/501100011033 and “ERDF A way of making Europe” and European Union NextGenerationEU/PRTR (“PID2022-143248OB-I00 and #CNS2022-135941) are acknowledged for funding. Additionally, the authors acknowledge the support of the European Union’s Horizon 2020 research and innovation program (grant agreement No 953169; **InterLynk**). DJdA thanks IKERBASQUE for sponsoring them.

