## Supporting Information for "Hybrid bioinks for embedded bioprinting of an artery model"

[1]U. Aizarna-Lopetegui, A. Herrero-Ruiz, K. Chavarri, M. Henriksen-Lacey, and D. Jimenez de Aberasturi.
CIC biomaGUNE, Basque Research and Technology Alliance (BRTA), 20014 Donostia-San Sebastián, Spain

[2]U. Aizarna-Lopetegui
Department of Applied Chemistry, University of the Basque Country, 20018 Donostia-San Sebastián, Gipuzkoa, Spain

[3] M. C. Decarli
University Medical Center Groningen (UMCG) / University of Groningen, 9713AV Groningen, The Netherlands

[4]L. Moroni
MERLN Institute for Technology-inspired Regenerative Medicine, Complex Tissue Regeneration Department, 6200MD, Maastricht, The Netherlands.

[5]M. Henriksen-Lacey
Centro de Investigación Biomédica en Red, Bioingeniería, Biomateriales y Nanomedicina (CIBER-BBN), 20014 Donostia-San Sebastián, Spain

[6]D. Jimenez de Aberasturi
Ikerbasque, Basque Foundation for Science, 48009 Bilbao, Spain

**Keywords**: embedded multilayered 3D bioprinting, stimuli-responsive inks, dECM-based bioinks, arteries, hybrid materials

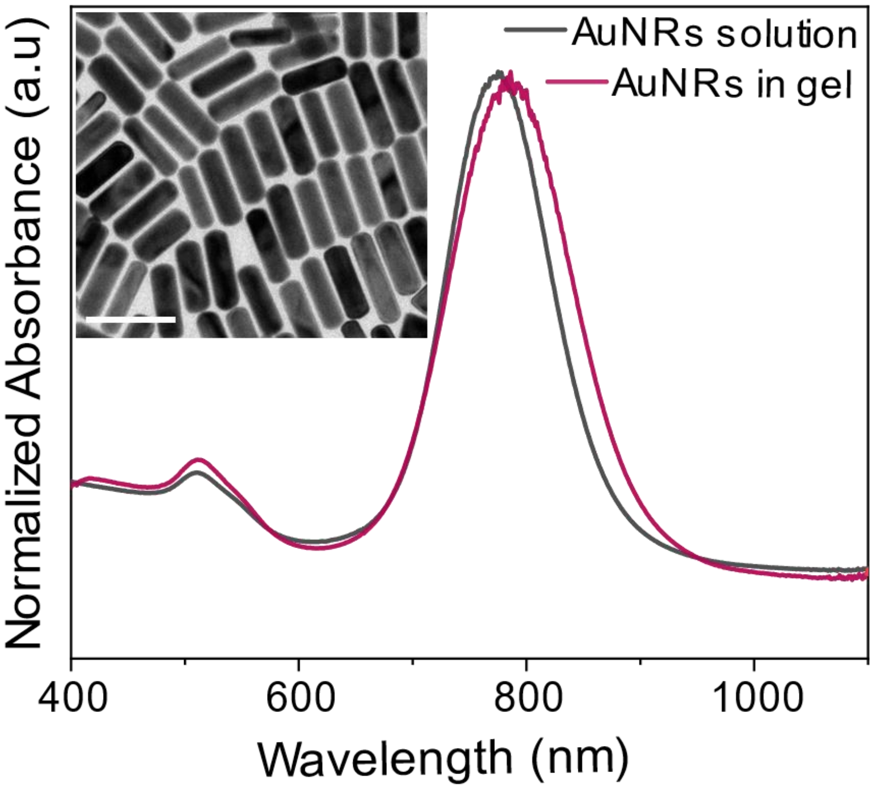


**Figure S1.** Characterization of AuNRs in solution and embedded in the hybrid gel synthetized in the microfluidic flow reactor by UV – vis spectroscopy (insert; scale bar 100 nm).

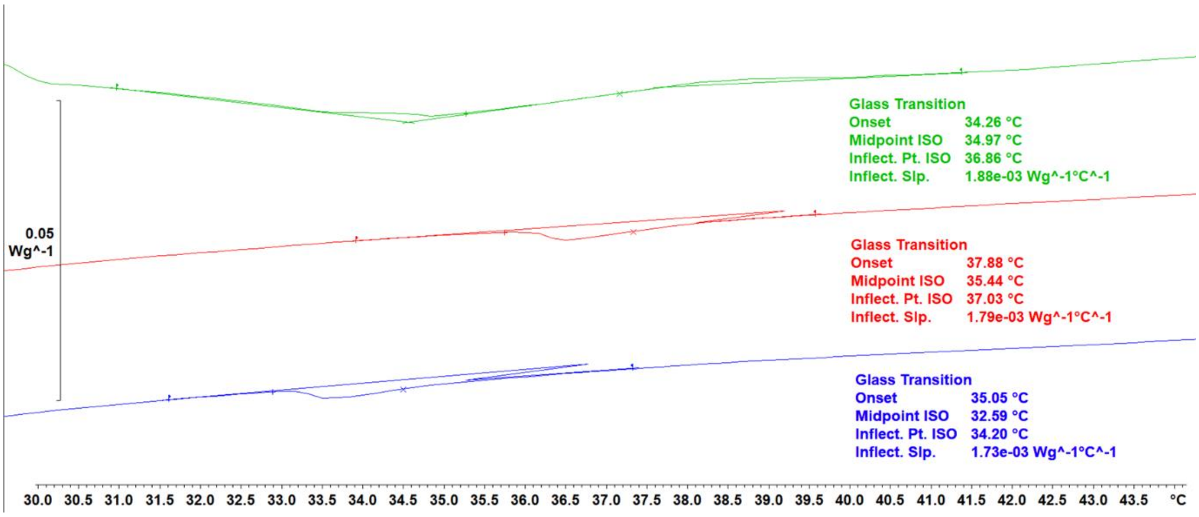


**Figure S2.** DSC curves of three heating cycles (10 - 50 °C at 5 °C/min) represented in blue (1st heating cycle), red (2nd heating cycle) and green (3rd heating cycle) of the final hybrid gel composition, employed to calculate the LCST (36 ºC) of the 0.5 mM AuNR-containing material through the mean of the infection points.

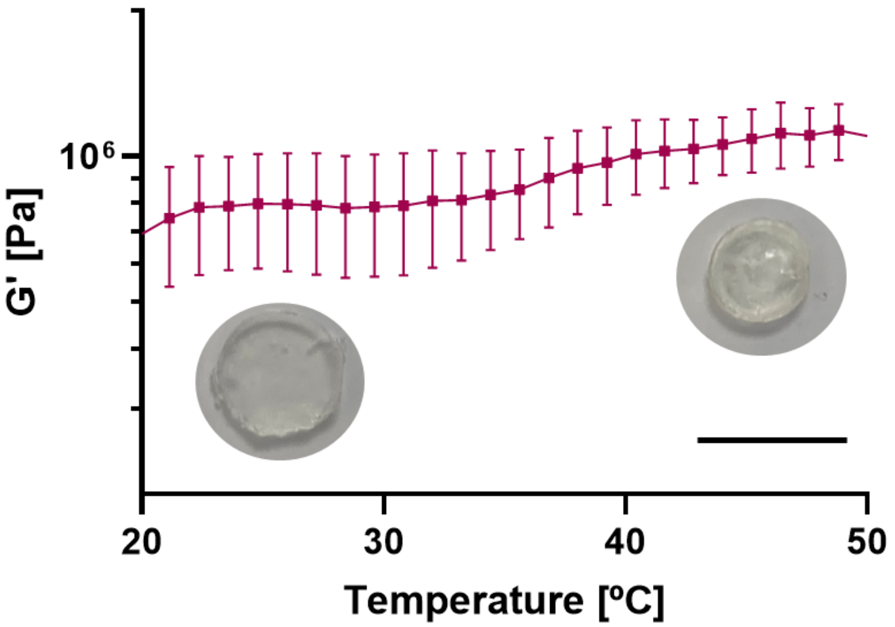


**Figure S3.** Temperature ramp (mean ± SD, n= 3) of final hybrid gel including heating from 20 °C (expanded gel) to 50 °C (contracted gel) at a heating rate of 1 °C/min (scale bar: 1cm).

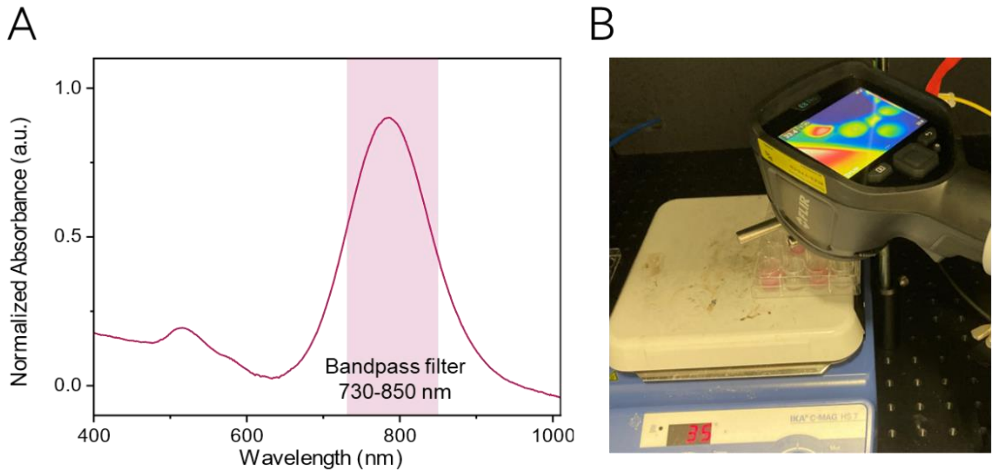


C

| Duty Cycle (%) | Minimum Temperature (ºC) | Maximum Temperature (ºC) | ΔT (ºC) |
|---|---|---|---|
| 15 | 35.0 | 36.9 | 1.9 |
| 17.5 | 35.7 | 37.6 | 2 |
| 20 | 35.9 | 39.2 | 3.3 |

**Figure S4.** A) UV–vis spectroscopy of AuNRs in solution and the bandwidth ranging from 730 to 850 nm, defined by the selected bandpass filter. B) Thermal characterization setup consisting of the laser, infrared camara, and heating plate. C) Changes in recorded temperature upon pulsed NIR irradiation with a fixed frequency of 10 Hz, at varying duty cycles.

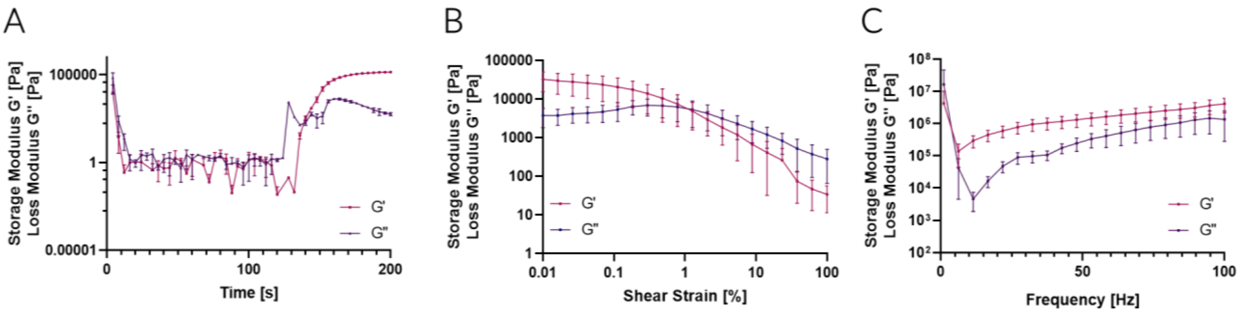


**Figure S5.** Rheological characterization of the viscoelastic properties of the hybrid ink, showing photo-crosslinking of the material upon (A) UV irradiation, (B) strain sweep, and (C) frequency sweep performed at 37 ºC (mean ± SD, n=3).

**Table S1.** Conditions for the decellularization of porcine pulmonary artery.

| Step | Time | Temperature |
|---|---|---|
| | Extract pulmonary arteries from porcine lung | |
| 0.5% (w/v) SDS – 0.5% (w/v) SDC | 72 h (90 min 0.9% (w/v) NaCl every 12 h) | RT |
| 0.9% (w/v) NaCl | 6 h | RT |
| 40 U/mL DNase in 1.5 mM $MgCl_2$ | O/N | 37 ºC |
| 0.9% (w/v) NaCl | 1 h 30 min | RT |
| 0.1% (v/v) PAA / 4% (v/v) EtOH | 2 h | RT |
| 2x PBS | 15 min | RT |
| 2x $dH_2O$ | 15 min | RT |

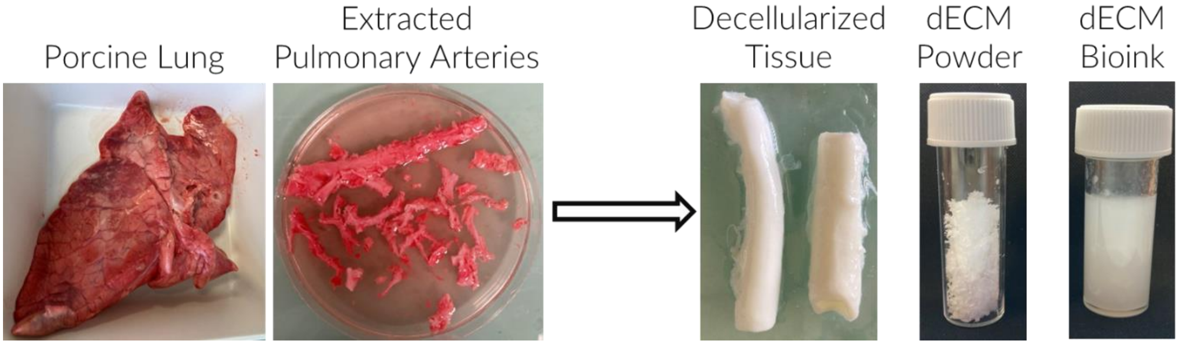


**Figure S6.** Macroscopic view of tissue obtained during the dECM bioink fabrication process.

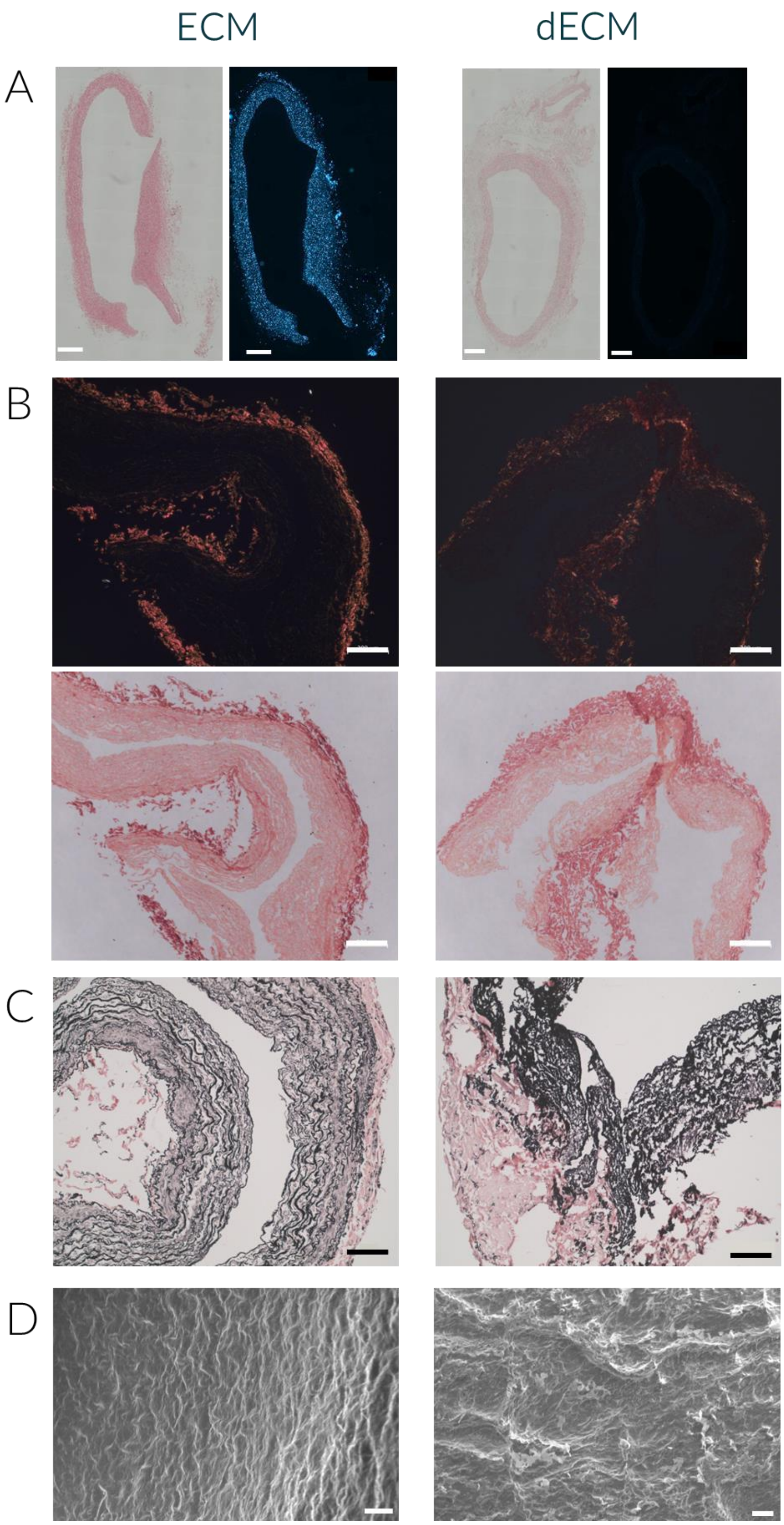


**Figure S7.** Histological characterization of the decellularization process including representative images of native (left) and decellularized (right) porcine pulmonary artery ECM.

A) H&E and DAPI staining (scale bars: 500 µm). B) Picrosirius red staining combined with polarized light (black and orange) and brightfield (white and red) imaging showing collagen fiber organization (scale bars: 200 µm). C) Verhoeff Van Giesson stain showing elastic fibers (black) (scale bars: 500 µm). D) SEM images of ECM and dECM transversal cuts (scale bars: 10 µm).

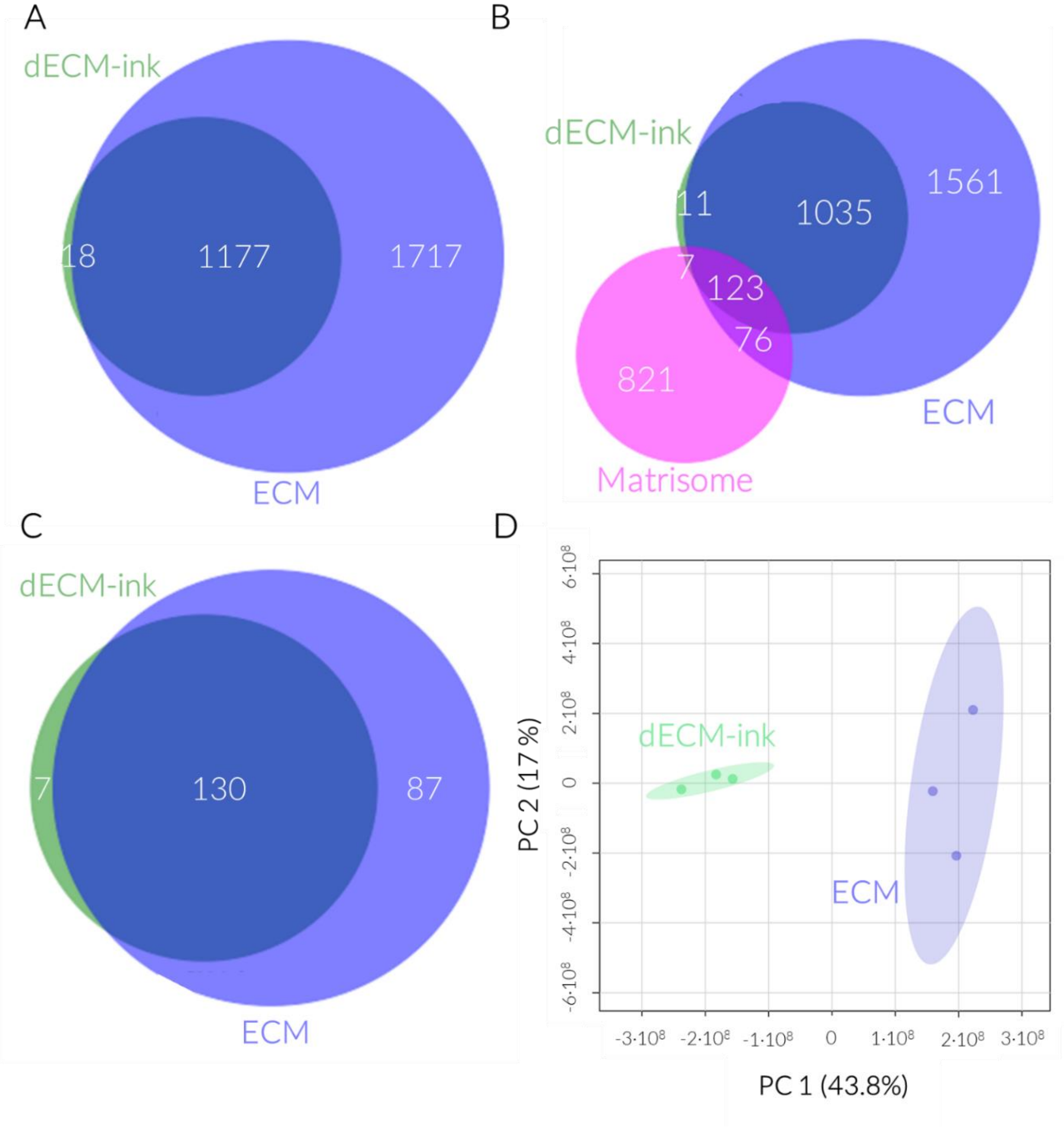


**Figure S8.** Venn diagrams illustrating the identified proteins in the ECM and dECM-ink showing (A) all identified proteins, (B) all identified genes with a focus on those described in the matrisome, and (C) proteins specifically associated with the matrisome. (D) PCA of the matrisome-associated proteins. The analysis included three biological replicates for each ECM and dECM-ink.

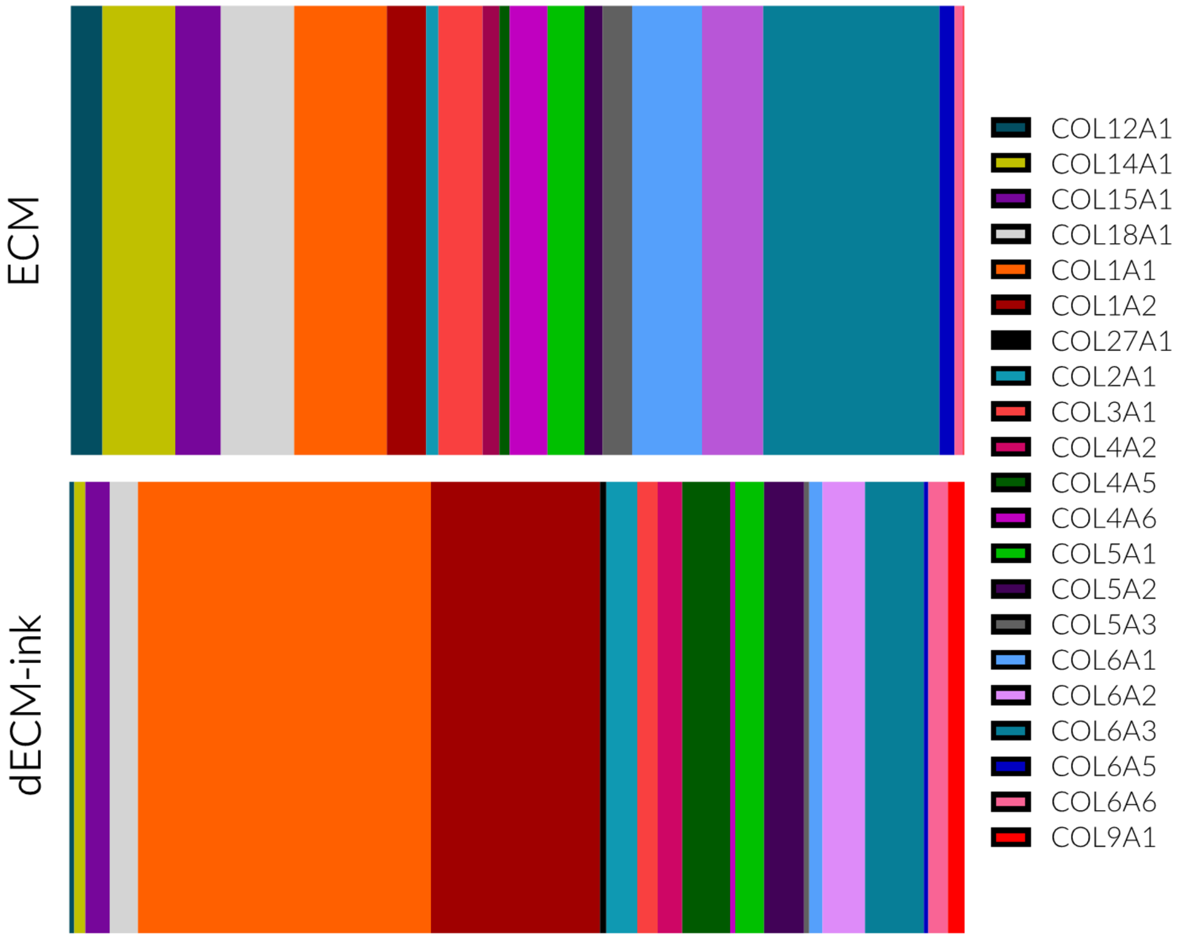


**Figure S9.** Relative intensities of the genes of the matrisomal profile of the ECM and the dECM-ink.

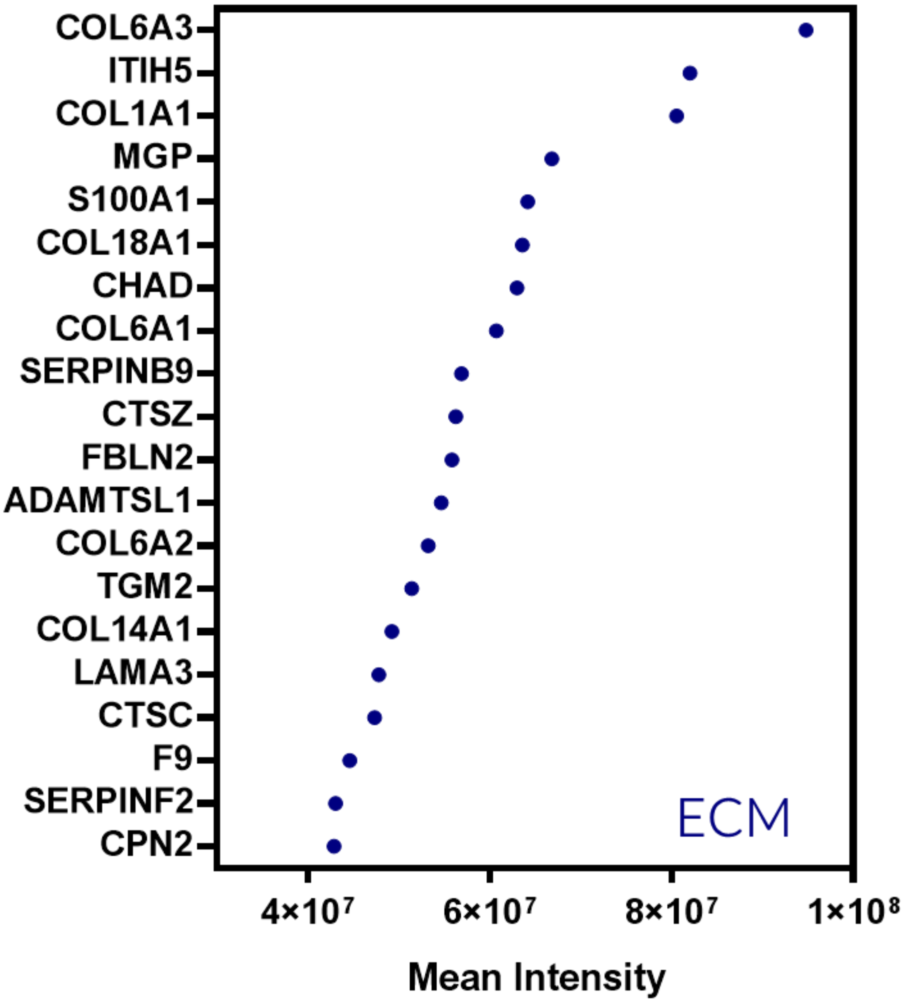


**Figure S10.** 20 most abundant genes associated to the porcine pulmonary artery ECM.

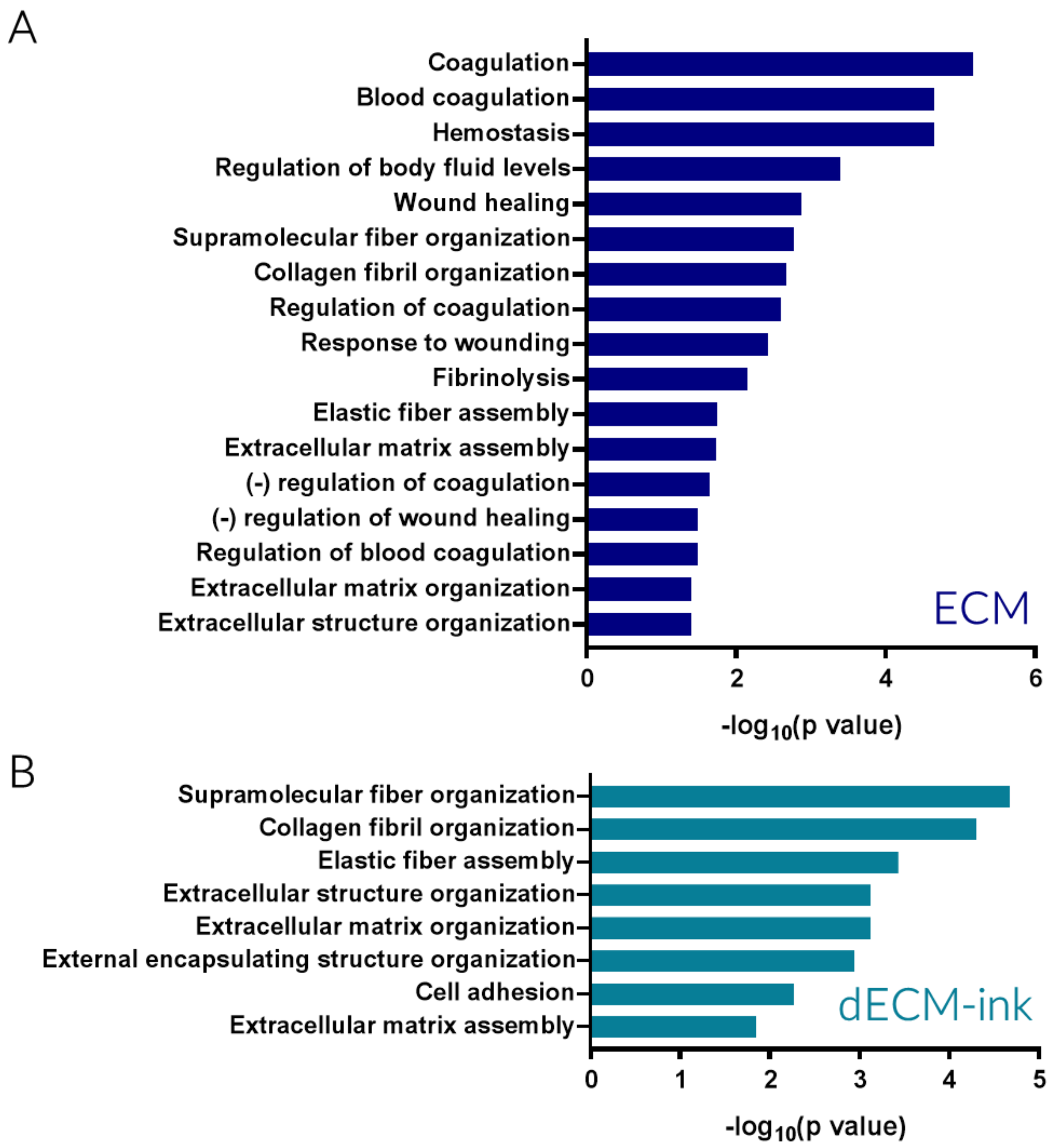


**Figure S11.** Gene ontology (biological process) of the complete set of identified genes in the (A) ECM and the (B) dECM-ink.

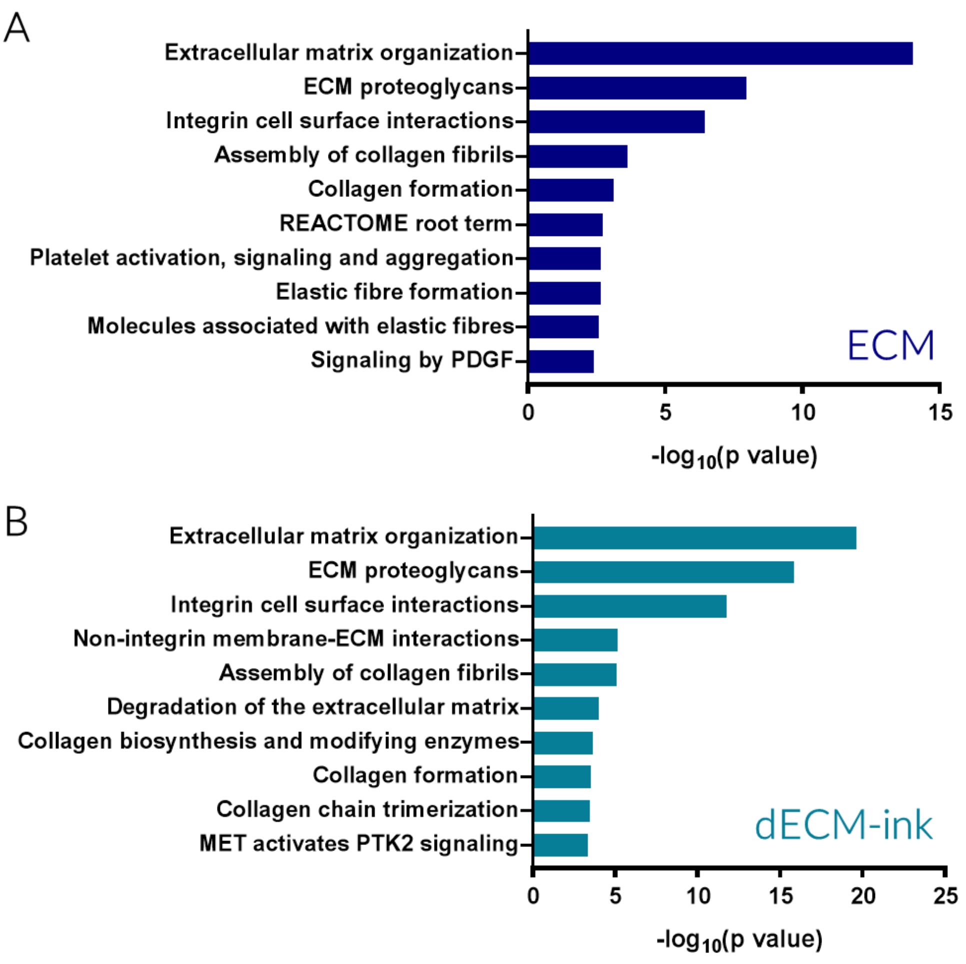


**Figure S12.** Reactome of the complete set of identified genes in (A) the ECM and (B) the dECM-ink.

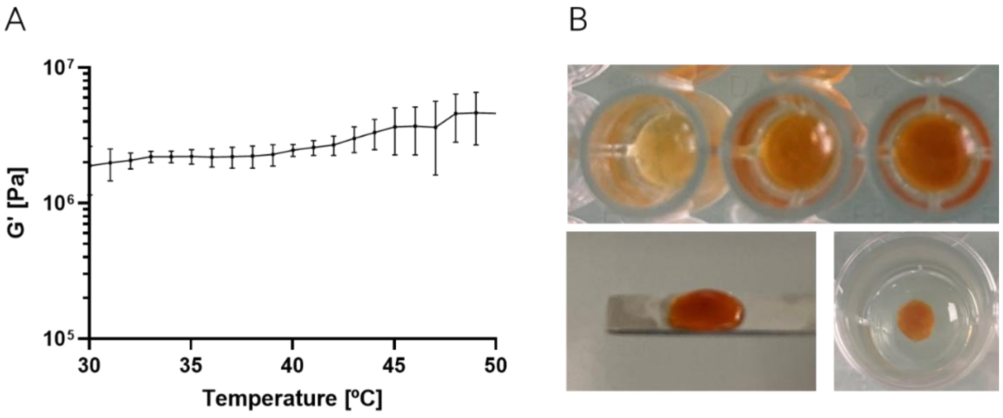


**Figure S13.** Characterization of the thermal and light-based crosslinking of the dECM. A) Temperature ramps with heating (from 30 to 50 ºC at 1 ºC/min) cycle (mean ± SD, n=3). B) Macroscopic images representing the Ru/SPS photo-polymerization mediated by 405 nm light irradiation, showing structural stability after crosslinking.

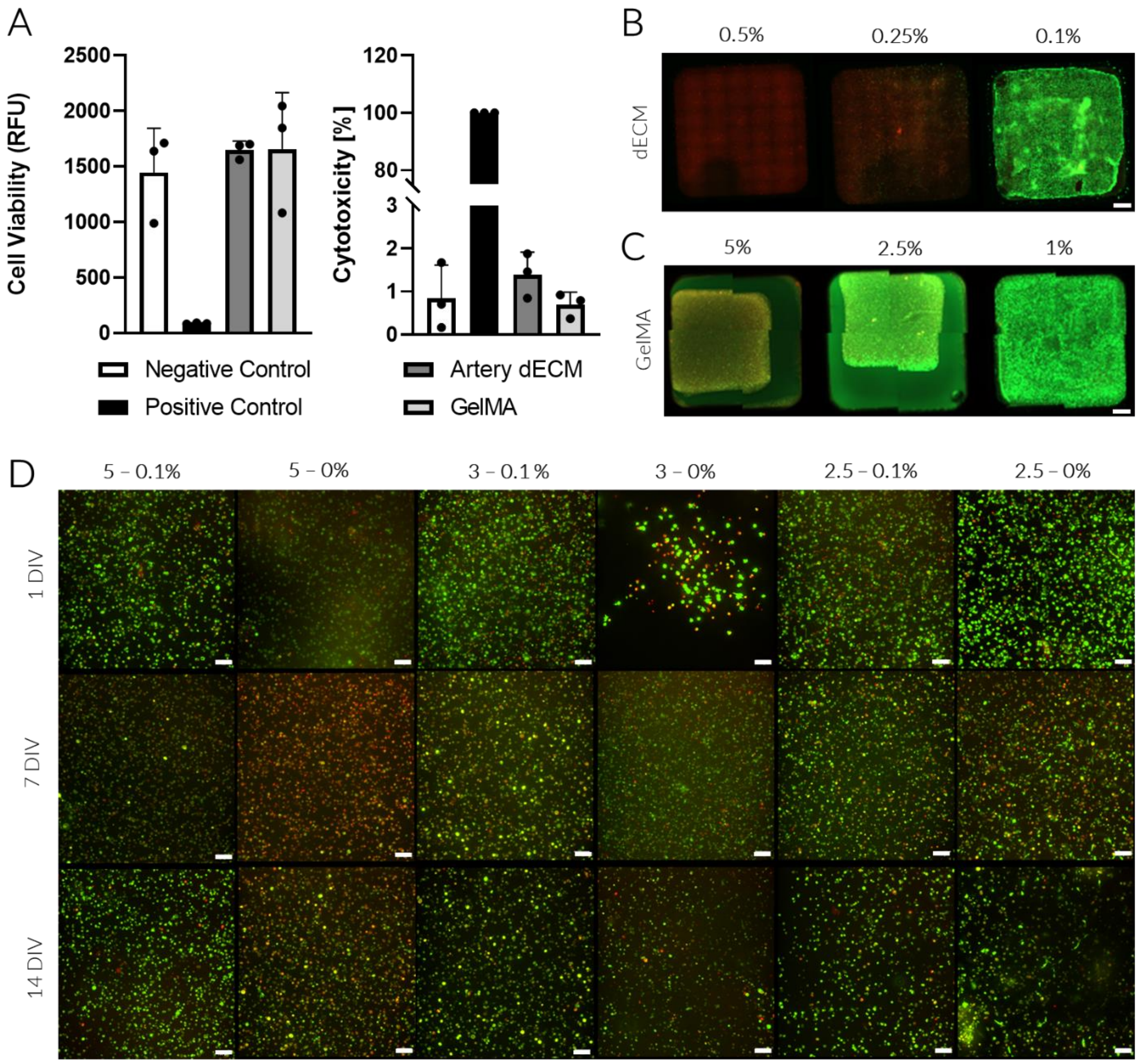


**Figure S14.** Evaluation of bioink biocompatibility with vSMCs. A) Bar plot showing cell viability and cytotoxicity for vSMCs, cultured in media but placed in contact with dECM and GelMA gels, assessed using Alamar Blue and LDH assays. B-C) Tiles reconstruction showing live/dead (green/red) fluorescence of vSMCs embedded in dECM or GelMA gels of various concentrations (scale bars: 1000 µm). D) Representative MIP images (zstack = 200 µm) showing live/dead (green/red) fluorescence of vSMCs embedded in dECM and GelMA gels of various concentrations. (scale bars: 100 µm).

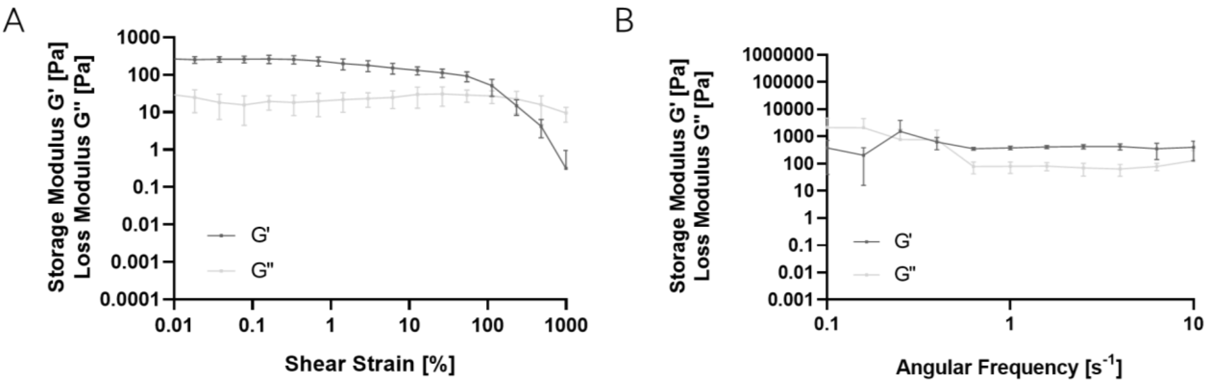


**Figure S15.** Strain and frequency sweeps of the 0.1 % (w/v) dECM and 3 % (w/v) GelMA bioink at 37 ºC.

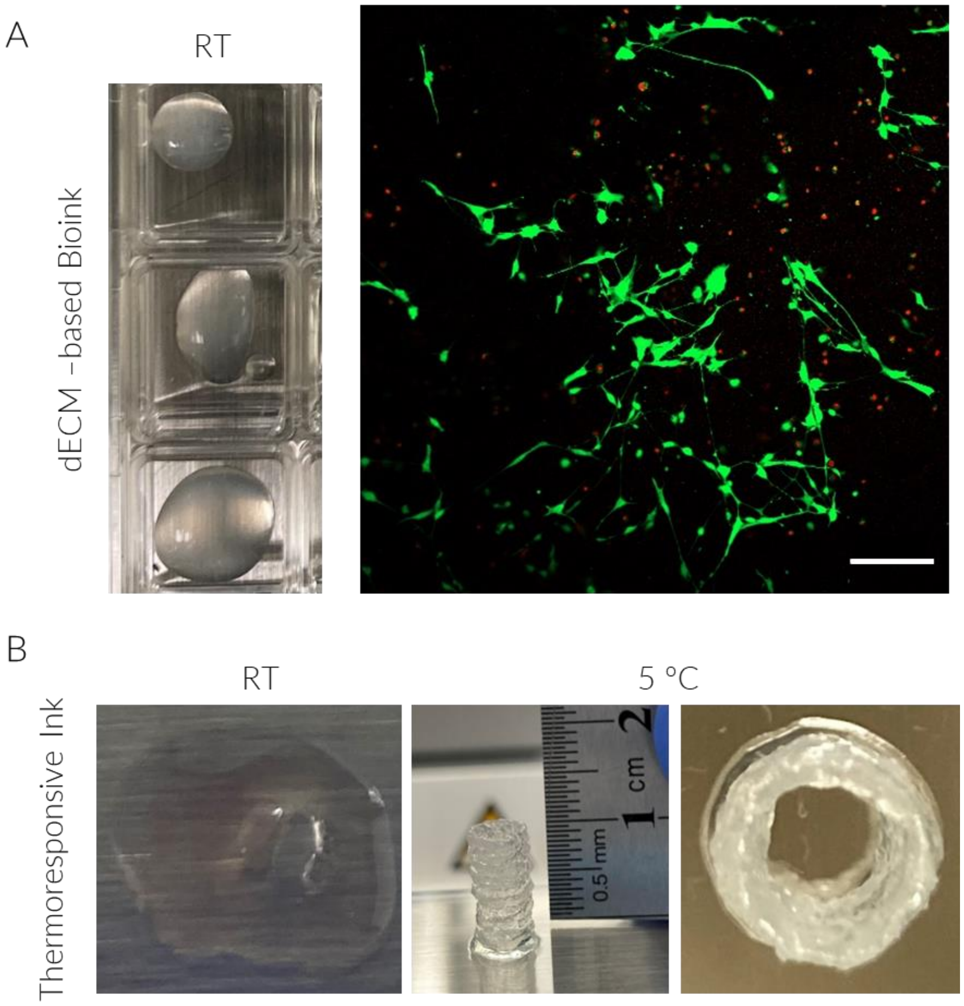


**Figure S16.** Extrusion-based printing of inks. A) Printing of the dECM-based bioink presenting low printing fidelity at RT and high post-printing cell viability and spreading (scale bar: 200 μm). B) Hybrid ink printed with low printing fidelity at RT, and high printing resolution at 5 ºC resulting in the fabrication of 1 cm high cylinders.

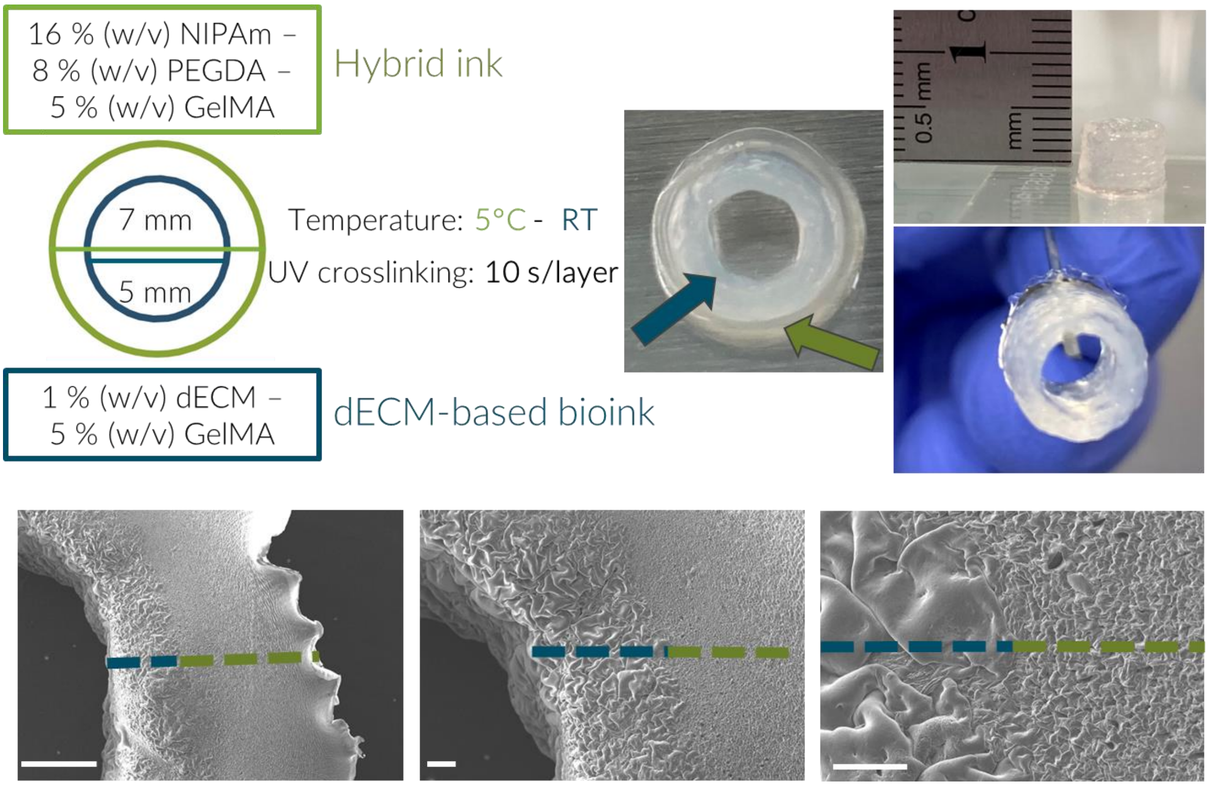


**Figure S17**. Extrusion-based 3D bioprinting of a concentric multilayered artery model including a representation of the macrostructure (top) and microstructure via SEM (bottom) (scale bars: 500 µm, 100 µm and 50 µm).

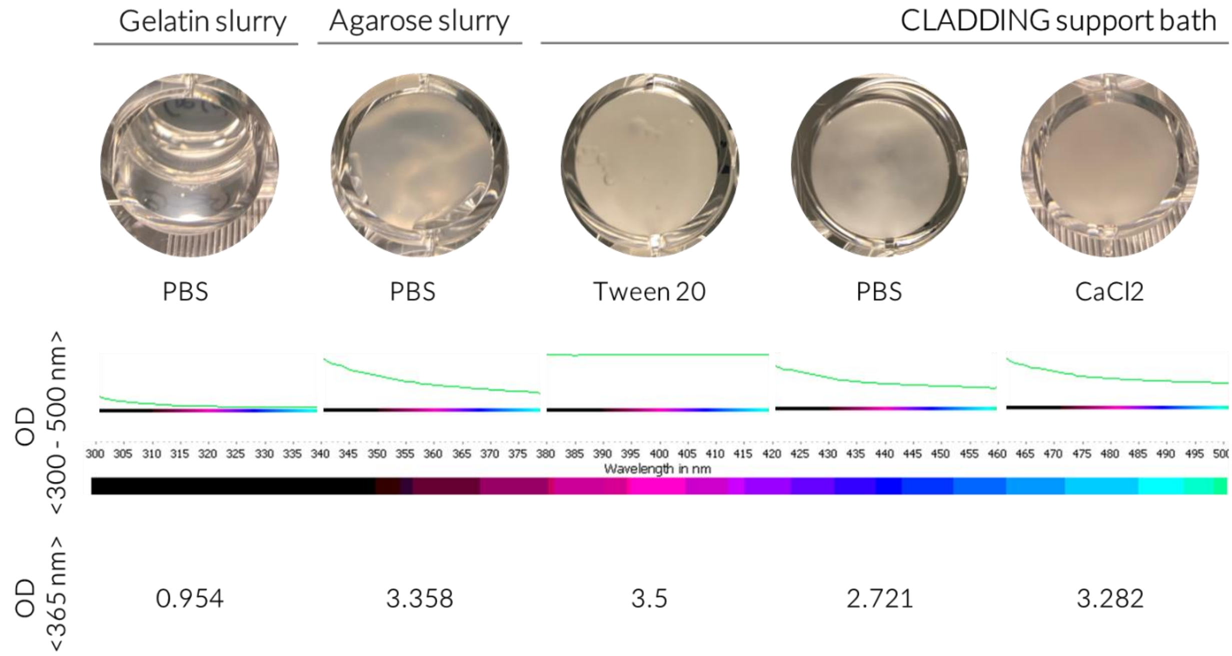


**Figure S18.** Characterization of the absorbance of the various slurry baths explored for assisted 3D printing including spectra showing the optical density at RT, over the range of 300 – 500 nm, and specifically at the wavelength employed for the photo-crosslinking (365 nm).

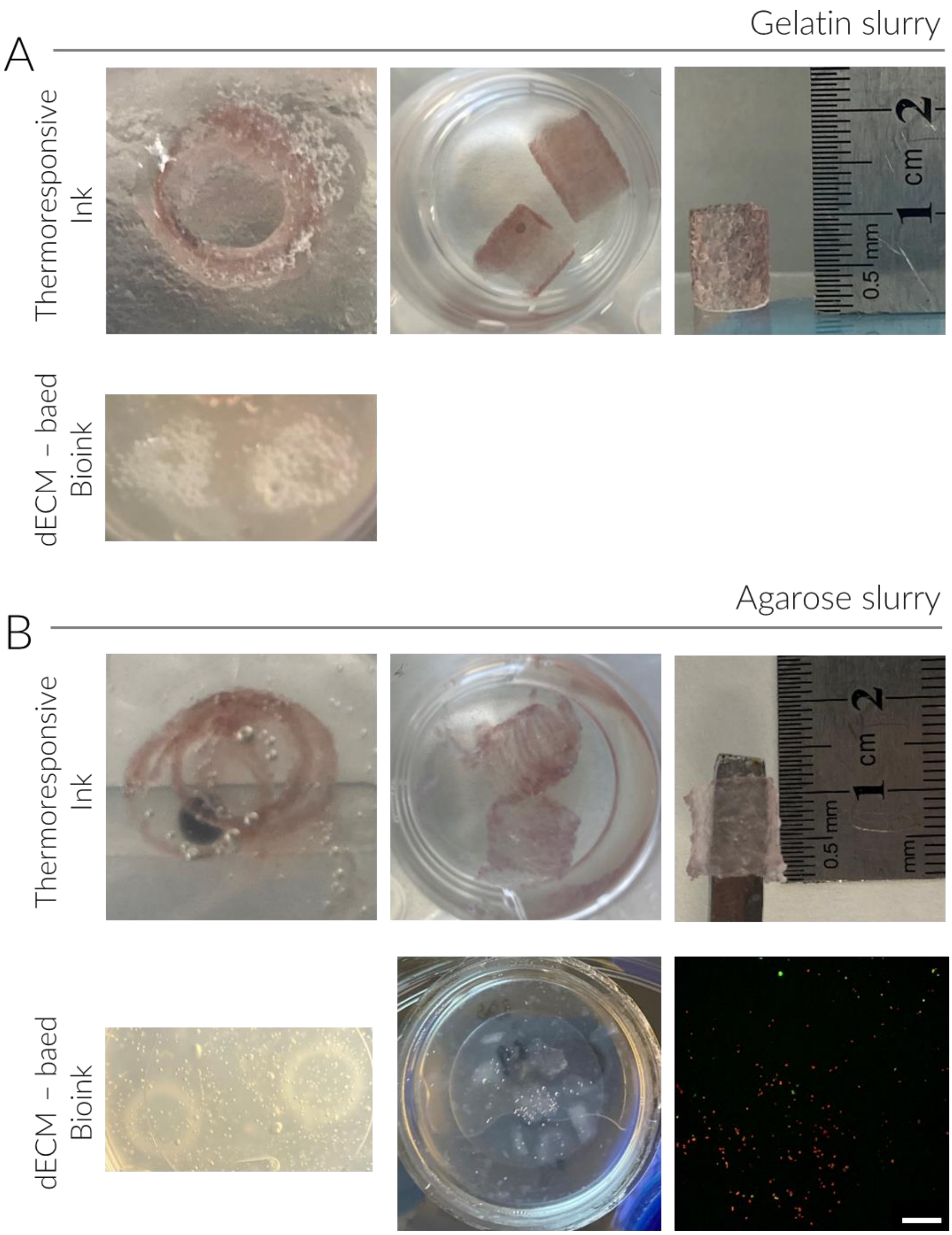


**Figure S19.** Example images of embedded 3D printing of the inks in gelatin (A) and agarose (B) slurry support baths. Live/dead fluorescence imaging of vSMCs embedded in the dECM-based bioink printed in agarose is shown at the bottom right (scale bar: 200 μm).

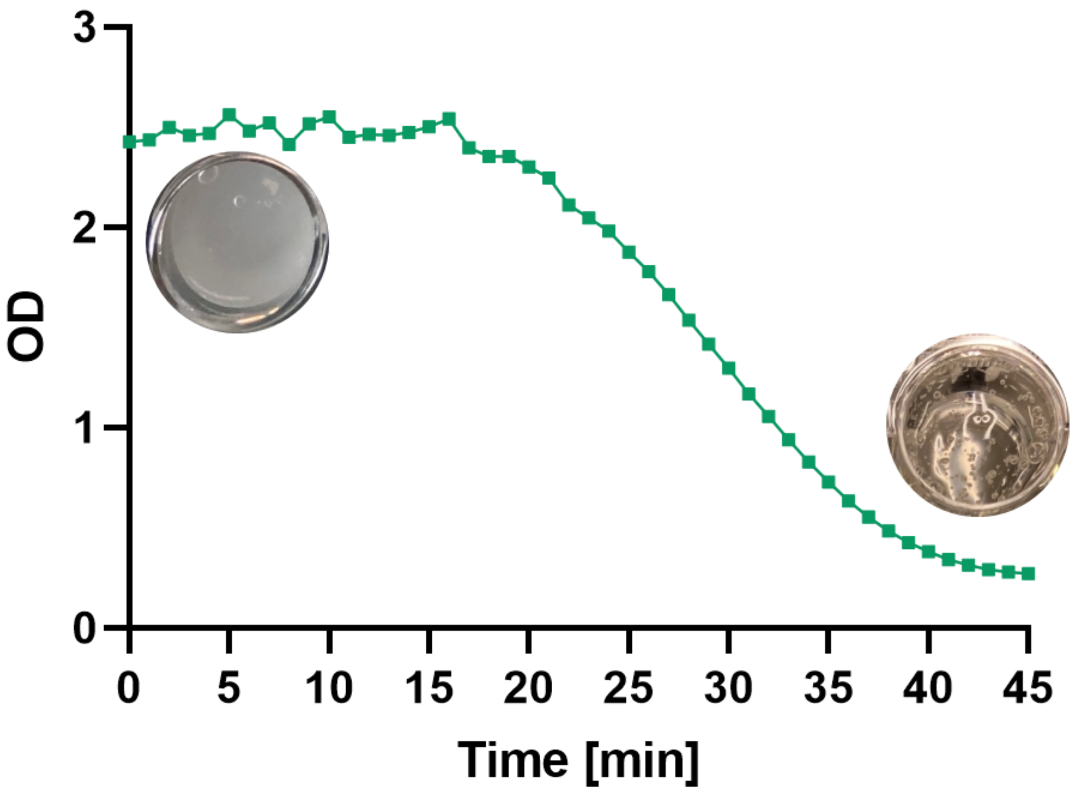


**Figure S20.** Changes in optical density (OD) of the *CLADDING* support bath at 37 ºC with time.

**Table S2.** Summary of the optimized parameters for embedded 3D bioprinting of the different materials.

| | **Hybrid ink** | **dECM-based Bioink** |
|---|---|---|
| Tip diameter (mm) | 0.2 – 0.3 mm | 0.2 – 0.3 mm |
| Tip length (mm) | 25.4 mm | 25.4 mm |
| Pressure (kPa) | 15 – 25 kPa | 15 – 25 kPa |
| Feed Rate (mm/s) | 15 mm/s | 10 mm/s |
| Layer Thickness (mm) | 0.2 – 0.3 mm | 0.2 – 0.3 mm |
| Angle between layers (º) | 60º | 60º |

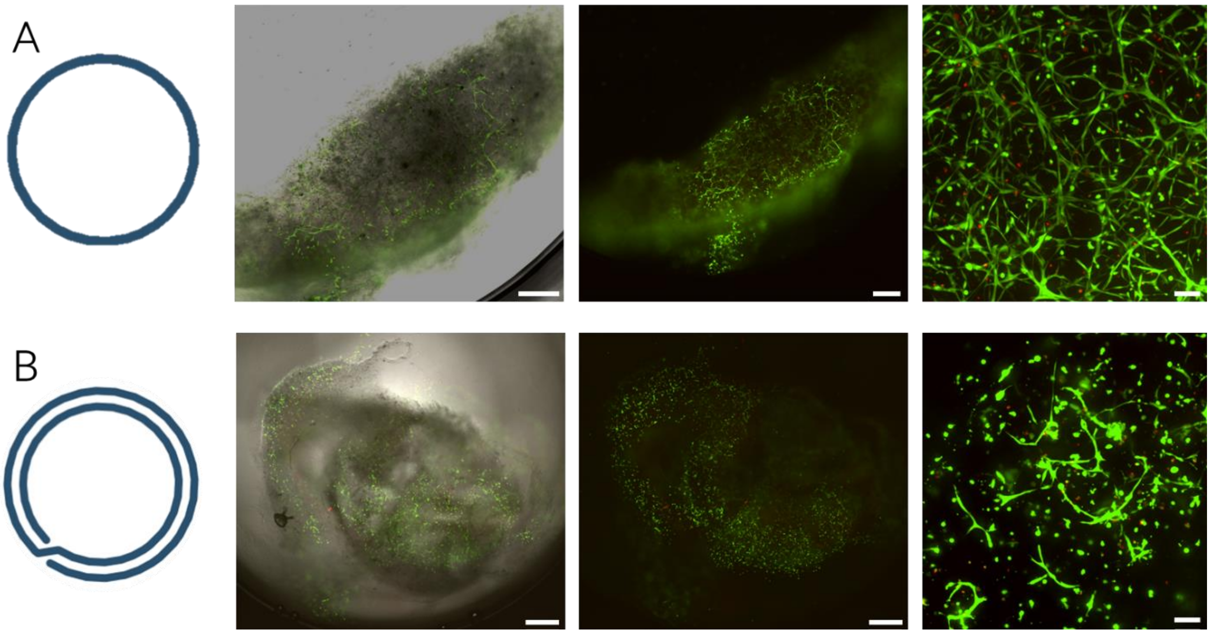


**Figure S21.** Embedded-printing of single (A) and doble-layer (B) cylinders of vSMCs in dECM-GelMA bioink represented by brightfield and fluorescence images of live/dead assay staining (scale bars: (A) left, 500 µm; middle, 500 µm; right, 100 µm, (B) left, 1000 µm; middle 1000 µm; right 100 µm).

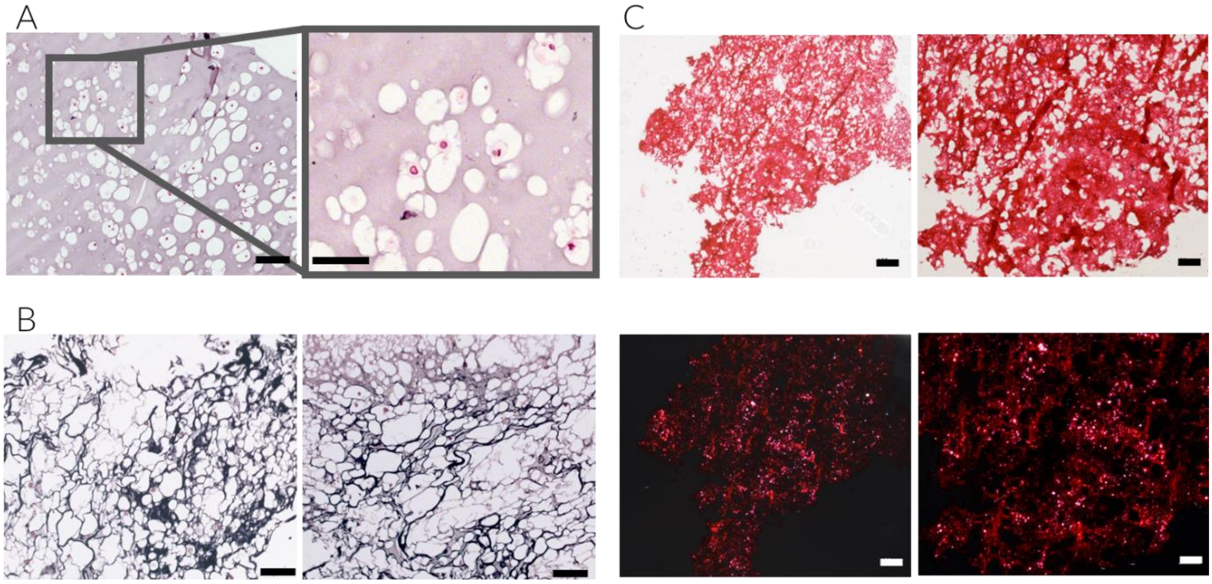


**Figure S22.** Histological evaluation of the embedded printed vSMC-based gels at day 7 including (A) representative images of H&E (scale bars: left; 100 µm, right; 50 µm), and analysis of the structure and composition through the study of the presence of (B) elastic fibers by Verhoeff Van Giesson stain (scale bars: 100 µm) and (C) collagenous fibers by brightfield images (top) and polarized light (bottom) of Picrosirius Red staining (scale bars: left; 100 µm, right; 50 µm).

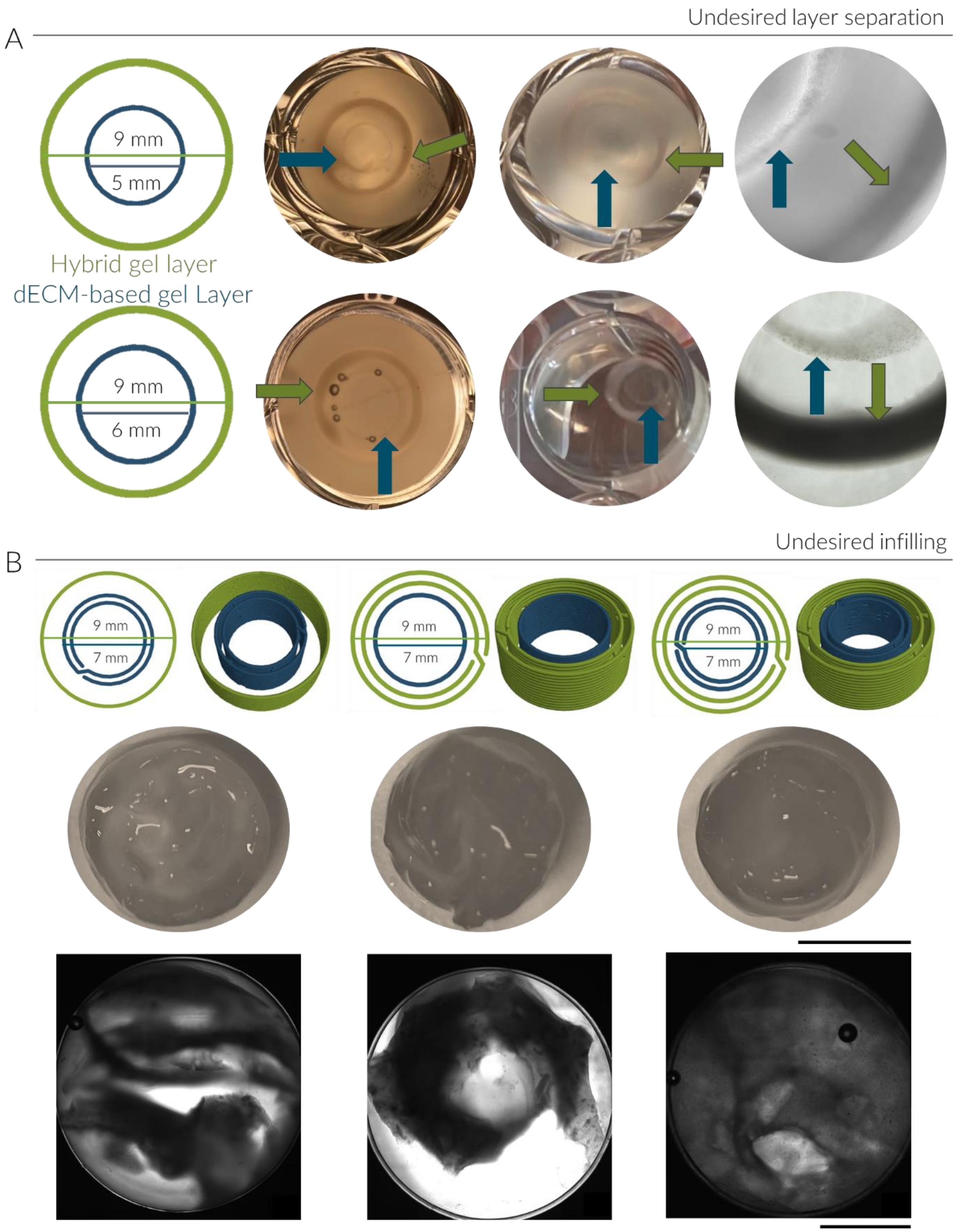


**Figure S23.** Optimization of the embedded 3D printing through the adjustment of the inner dECM-layer diameter (A) and number of layers of the model (B) (scale bars: 0.5 cm).

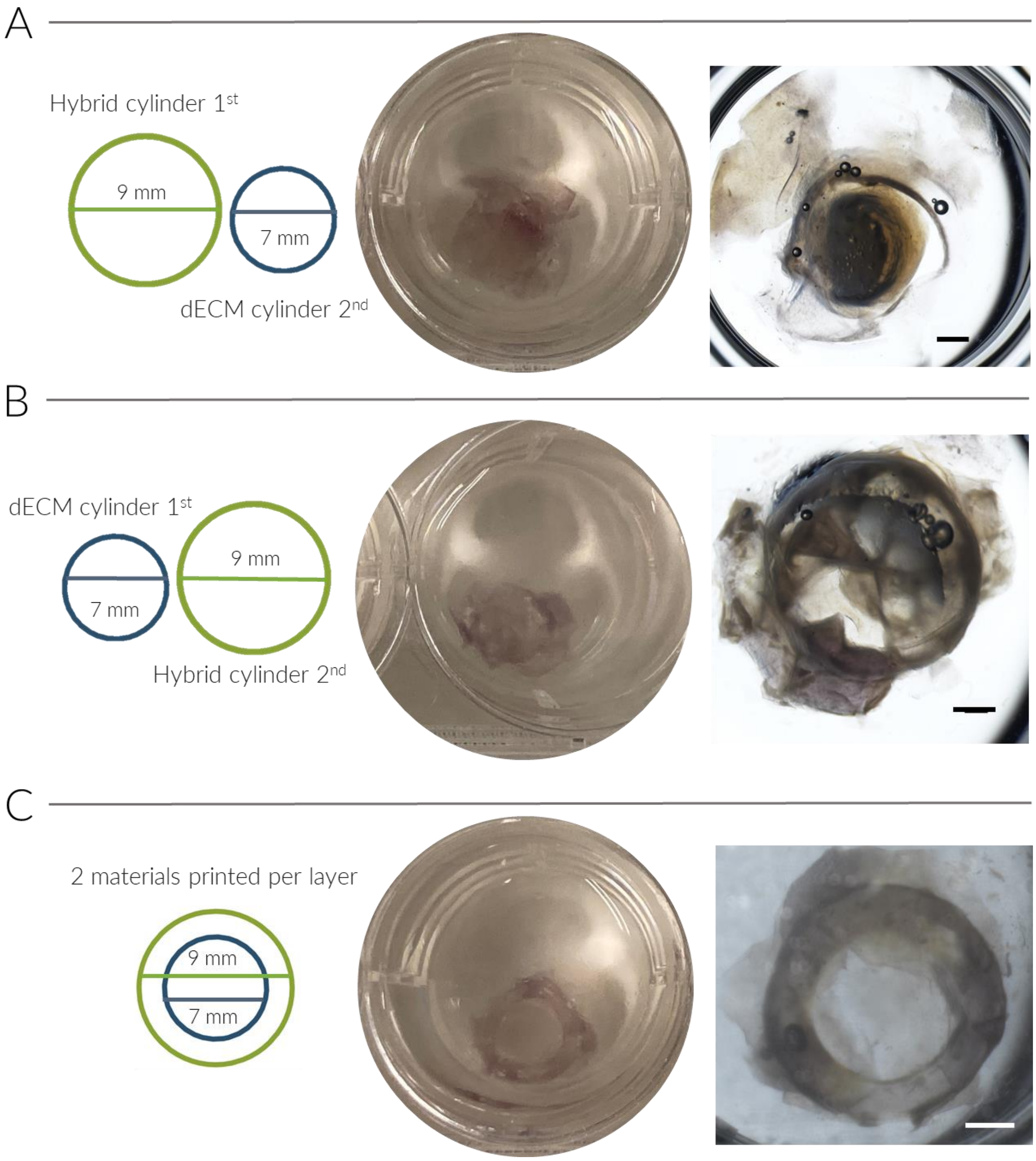


**Figure S24.** Optimization of the printing sequence for fabricating concentric multi-material cylinders involved testing different approaches: (A) continuous printing of the hybrid outer cylinder followed by the dECM-based bioink as the inner layer, (B) printing the complete inner dECM-based layer first, followed by the external hybrid layer, and (C) an optimized sequence where both materials are printed layer by layer simultaneously (scale bars: 2cm).

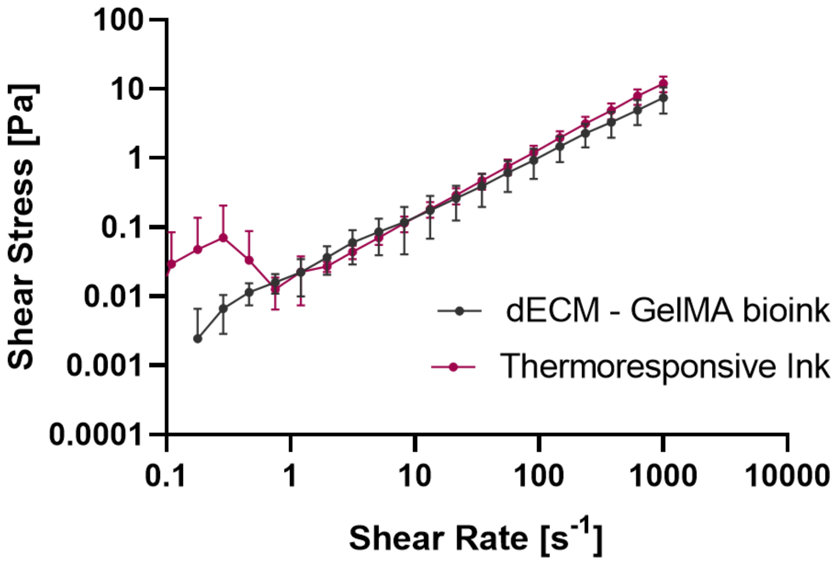


**Figure S25.** Flow curves of both inks at varying shear rates (mean ± SD, n=3).

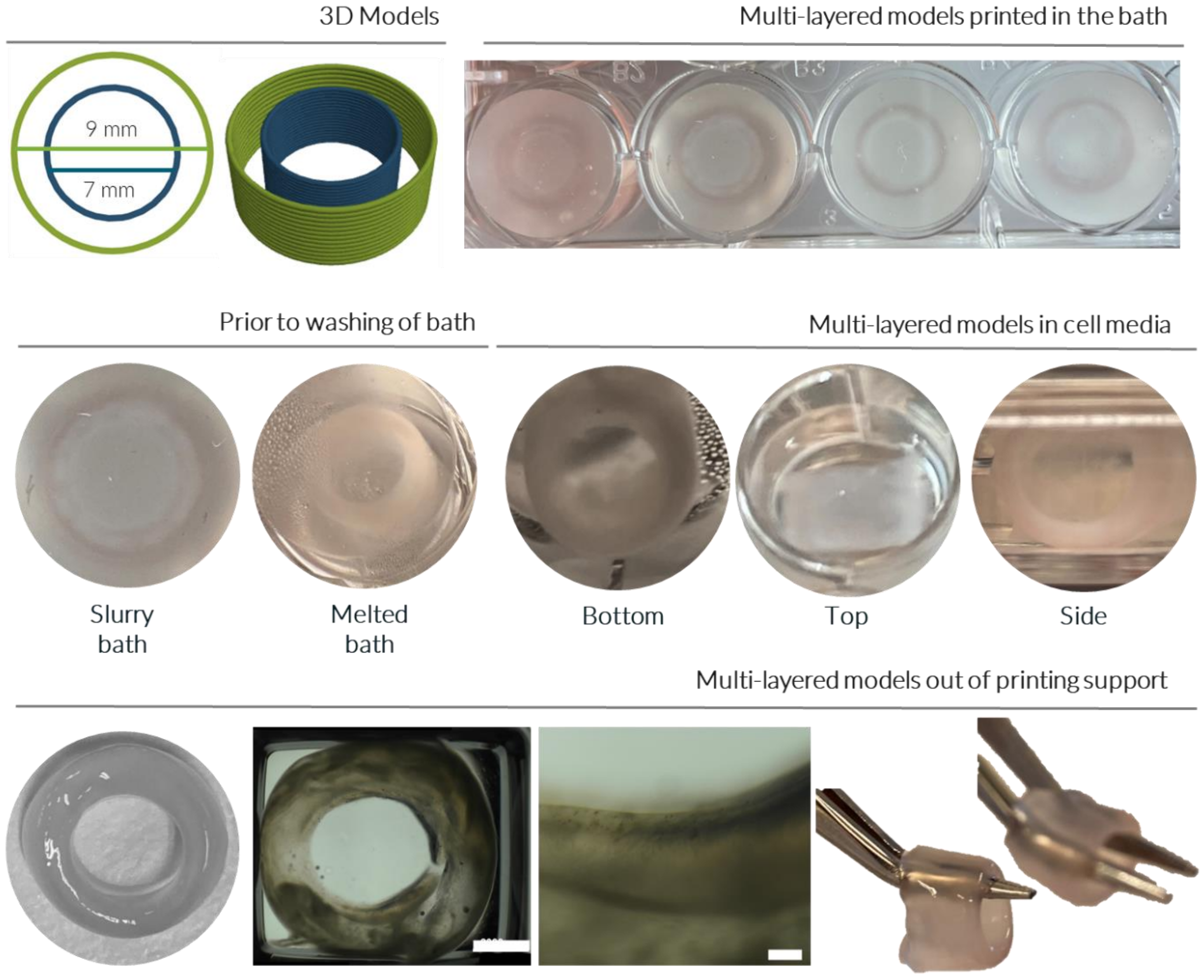


**Figure S26.** Examples of multi-material concentric cylinders produced using embedded 3D printing are illustrated at each step. Macroscopic images depict the constructs throughout the

3D printing process, encompassing the design of the 3D models and demonstrating the high reproducibility of printing within the support bath. These multi-layered models exhibit structural integrity both within the slurry support bath and after melting, and they maintain this stability upon removal and immersion in cell media. The high structural integrity allows these constructs to be transferred out of the printing support and manipulated easily without breakage, facilitating persistent characterization efforts (scale bars: 2 cm and 200 μm).

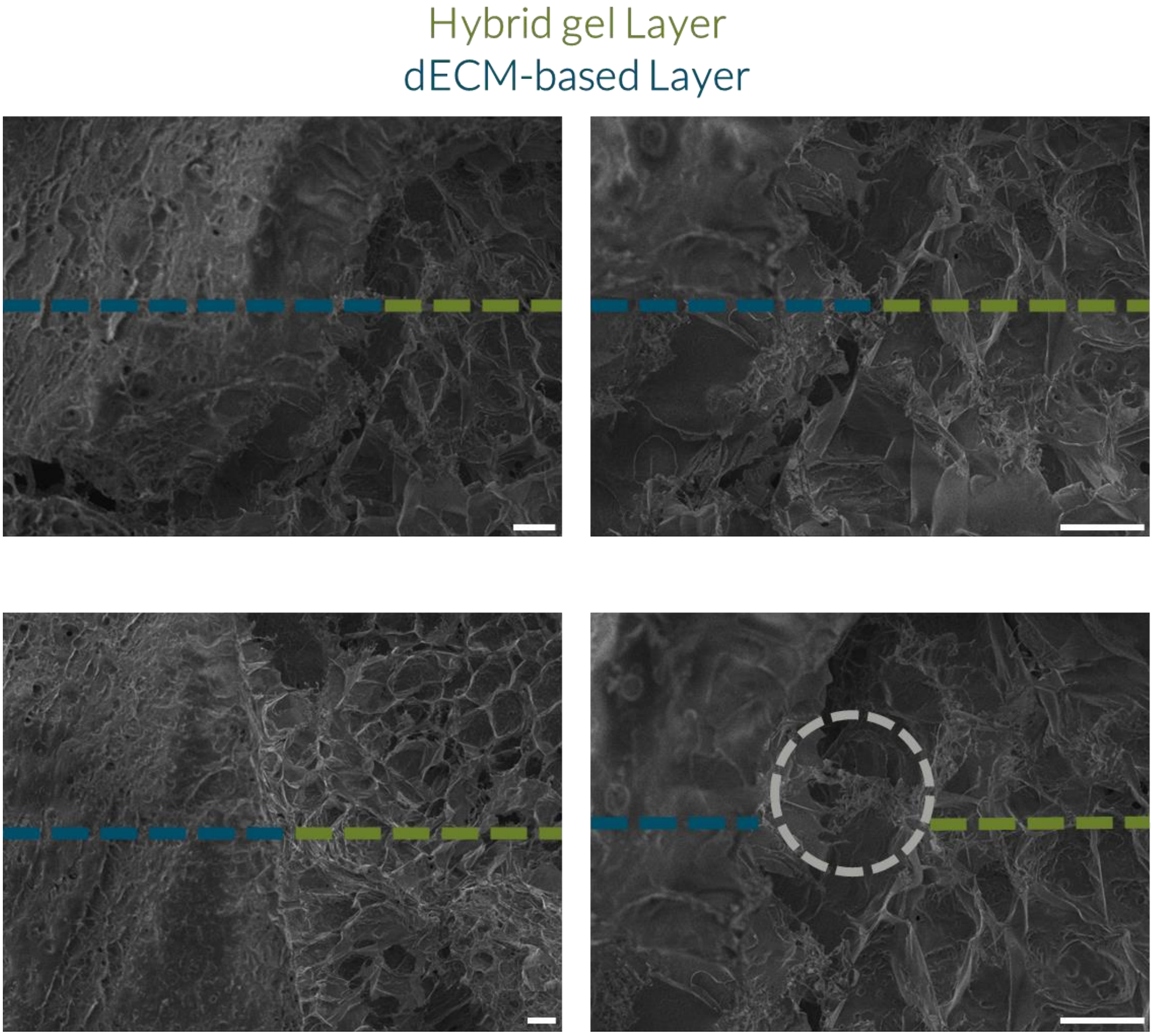


**Figure S27.** SEM images of the interface of the hybrid gel and dECM-based layers of the 3D printed multi-material constructs at different magnification (scale bars: 100 μm).

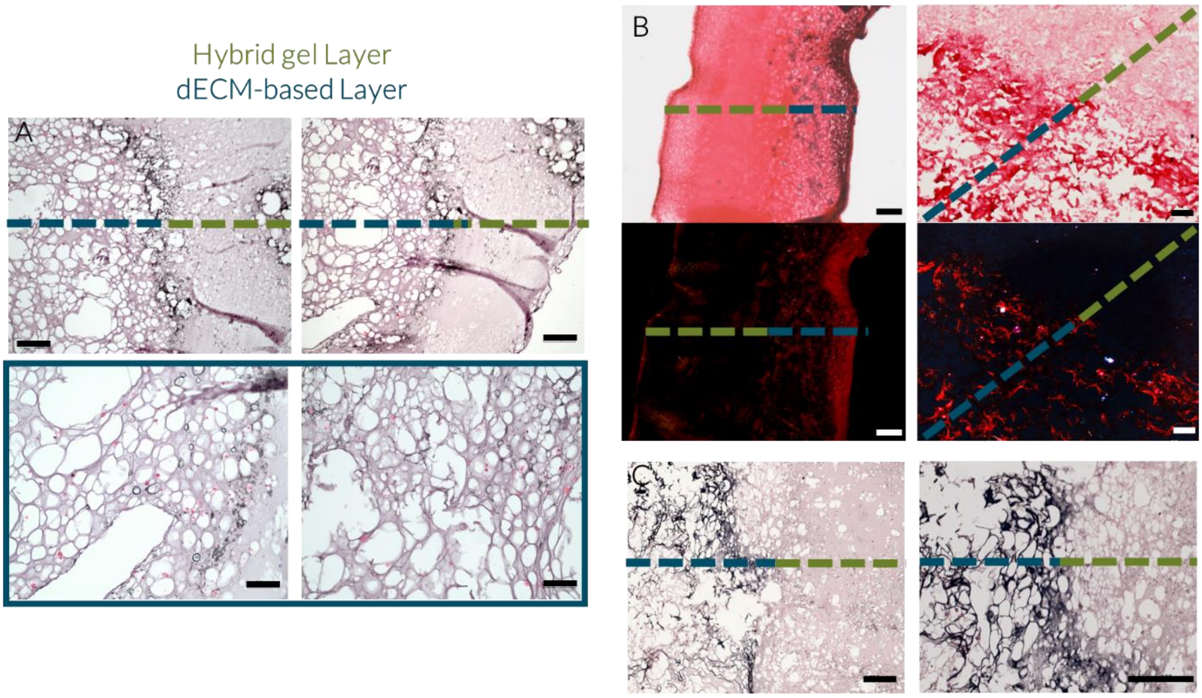


**Figure S28**. **(A)** Representative images of H/E of multilayered constructs (top) and vSMC-based layer (bottom) (scale bars: top; 200 µm, bottom; 100 µm). Analysis of the structure and composition of the multilayered structures printed within the CLADDING support bath through the study of the presence of **(B)** collagenous fibers by brightfield images (top) and polarized light (bottom) of Picrosirius Red staining (scale bars: left; 100 µm, right; 50 µm) and **(C)** Verhoeff Van Giesson stain showing elastic fibers (scale bars: 200 µm).

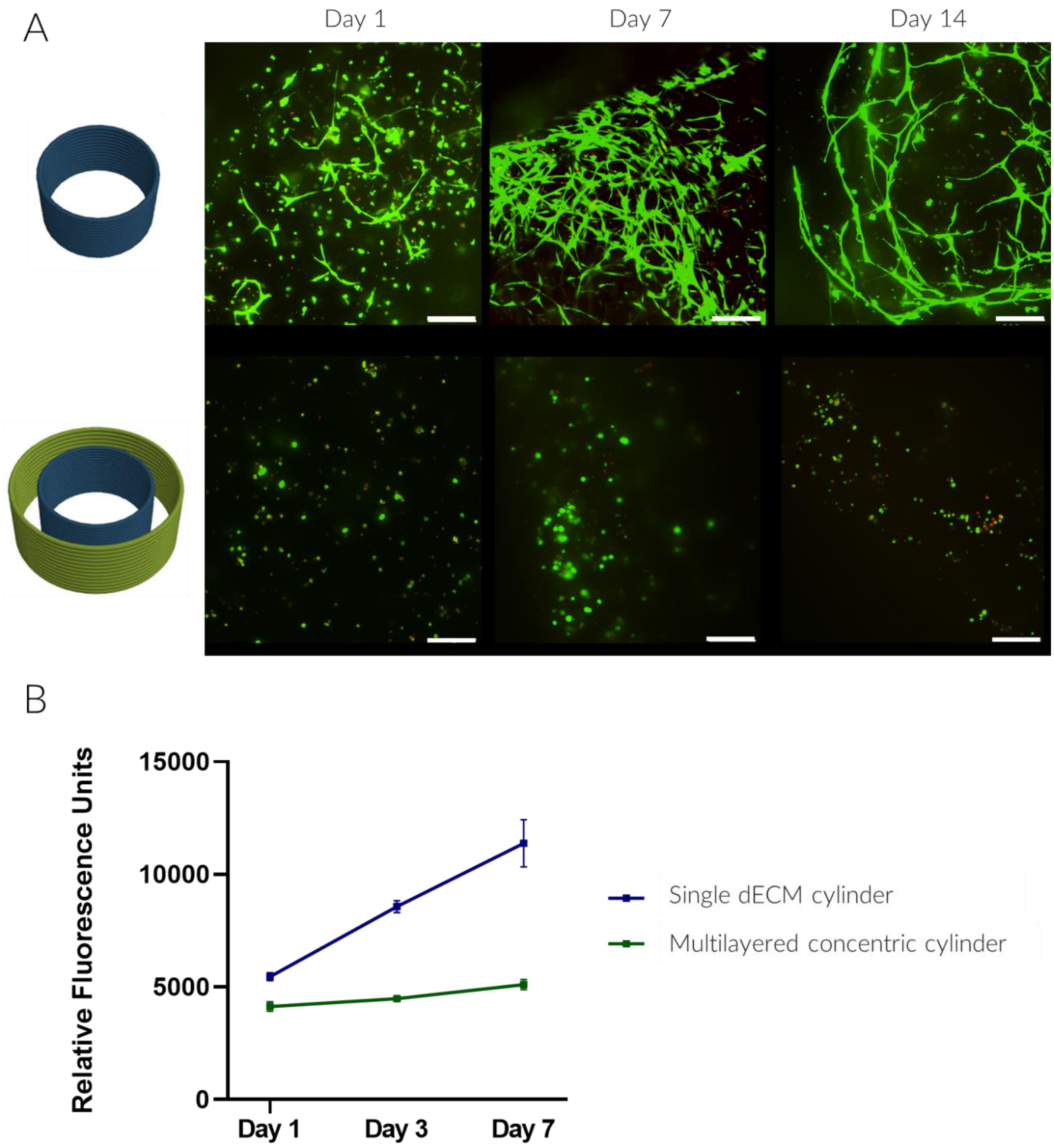


**Figure S29.** A) Characterization of 3D bioprinted constructs overtime via live/dead cell fluorescent imaging of single (top) and multilayered (bottom) cylinders (sacale bars: 200 µm). B) Cell metabolic activity measured using the PrestoBlue assay, of single (blue) and multimaterial (green) cylinders.

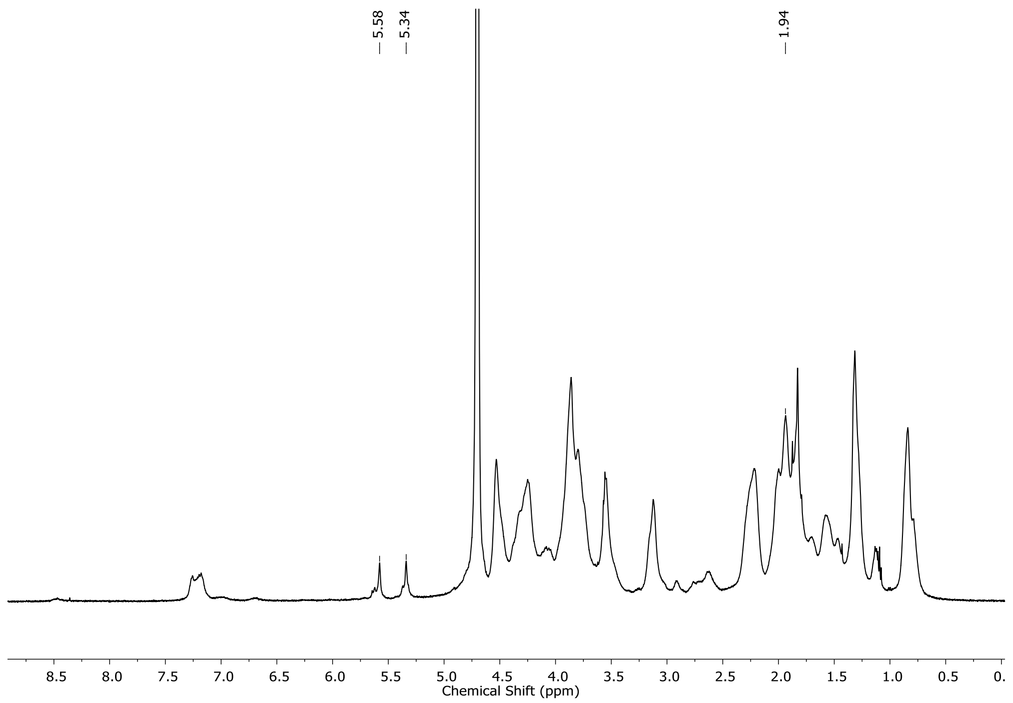


**Figure S30**. H-NMR spectra of GelMA.

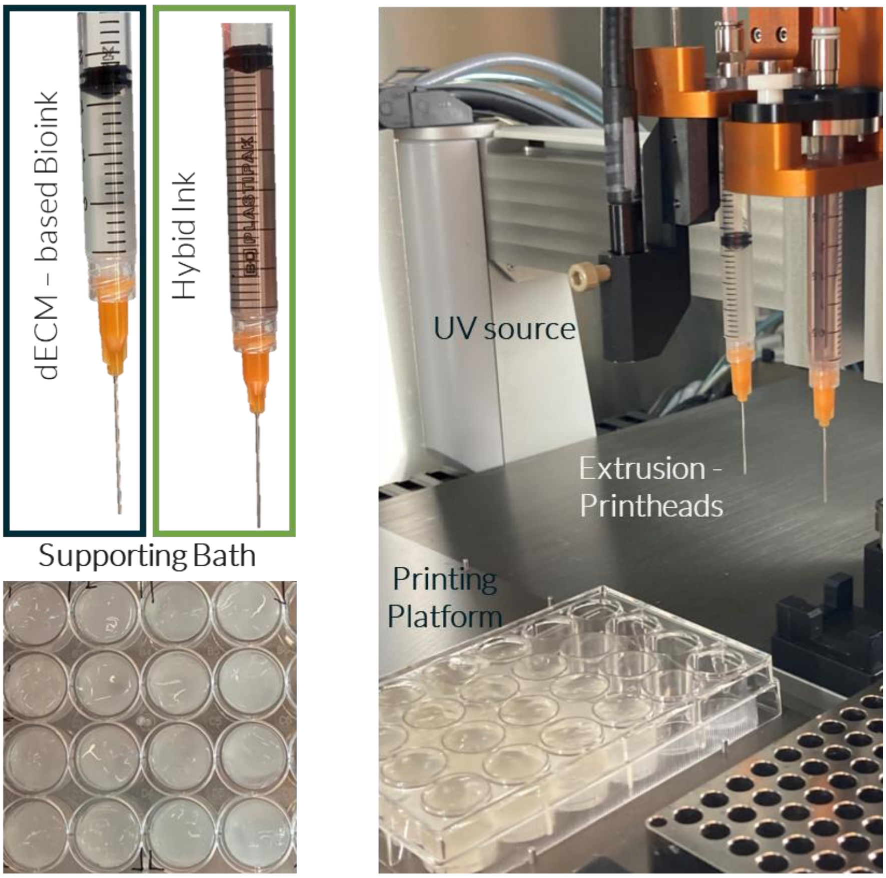


**Figure S31.** Bioprinting components and setup employed for embedded bioprinting of multi-material models.
